# Clearing the air on pollutant disruptions of the gut-brain axis: Developmental exposure to Benzo[a]pyrene disturbs zebrafish behavior and the gut microbiome in adults and subsequent generations

**DOI:** 10.1101/2024.08.01.606028

**Authors:** Alexandra Alexiev, Ebony Stretch, Kristin D. Kasschau, Lindsay B. Wilson, Lisa Truong, Robyn L. Tanguay, Thomas J. Sharpton

## Abstract

Developmental exposure to benzo[a]pyrene (BaP), a ubiquitous environmental pollutant, has been linked to various toxic effects, including neurodegenerative disorders and, most recently, multigenerational behavioral impairment. While the specific mechanisms driving BaP neurotoxicity are not fully understood, recent work highlights two important determinants of developmental BaP neurotoxicity: (1) the aryl hydrocarbon receptor (AHR), which is responsible for inducing host metabolism of BaP, and (2) the gut microbiome, which may interact with BaP to affect its metabolism, or be perturbed directly by BaP to yield disruptions to the gut-brain axis. To explore the role of AHR, the gut microbiome, and their interaction on BaP-induced neurotoxicity, we utilized the zebrafish model. We sought to determine (1) how exposure to BaP and developmental expression of AHR2, a key gene in the zebrafish AHR pathway, link to adult zebrafish behavior, (2) how these same variables associated with the structure and function of the adult zebrafish gut metagenome, and (3) whether these associations were multigenerational. This finding revealed a reticulated axis of association between BaP exposure, developmental AHR2 expression, the zebrafish gut metagenome, and behavior. Our results also indicate that AHR2 is a key mediator of how BaP elicits neurotoxicity and microbiome dysbiosis. Additionally, this axis of association manifests inter- and transgenerationally, suggesting that exposure to BaP may yield dysbiotic and neurodevelopmental impacts on subsequent generations. These findings demonstrate the power of utilizing the zebrafish model to study pollutant-metagenome interactions and elucidate the role of specific host genes in the neurotoxicity and dysbiosis.

**Importance:** Benzo[a]pyrene (BaP) is a toxic chemical that is especially ubiquitous in industrialized nations, as it is produced in large quantities by burning coal, oil, and other organic compounds. Early-life and *in utero* exposure to this chemical can induce adverse behavior changes in model animals, like mice and zebrafish. In humans, developmental exposure is linked to symptoms of ADHD, anxiety, and depression. This study found BaP affects zebrafish behavior and the gut microbiome throughout their life and across subsequent generations. We further discovered that a host regulatory pathway called the aryl hydrocarbon receptor is a key component of how benzo[a]pyrene, the gut microbiome, and behavior interconnect. The aryl hydrocarbon receptor binds many toxicants, as well as microbial metabolites associated with maintaining gut homeostasis, and regulates the gut-brain connection. This receptor presents a new potential mechanism of how gut microbiota relate to BaP exposure and behavior for future studies to investigate.

## Introduction

Benzo[a]pyrene (BaP) is a model environmental chemical toxicant that is often used to study the effect of chemical exposure on host health due to its ubiquity, impact on the early-life physiological and behavioral development of vertebrates, and ability to elicit such effects across multiple generations (Gelboin 1980; Bukowska, Mokra, and Michałowicz 2022; Knecht et al. 2017a; 2017b). Well-known as a pro-carcinogenic compound, studies have linked BaP to neurodevelopmental and neurodegenerative disorders (Perera et al. 2012; W. Zhang et al. 2016; Chepelev et al. 2015; Gao et al. 2017; Slotkin et al. 2019; Liu et al. 2020; Li et al. 2018). Experiments in vertebrate model systems, notably zebrafish, support these observations, finding that developmental BaP exposure drives neurodegenerative phenotypes (Gao et al. 2018; Knecht et al. 2017b; 2017a; Perera et al. 2012; W. Zhang et al. 2016). These neurotoxic effects also appear to manifest in subsequent, unexposed generations (Knecht et al. 2017b; Pandelides et al. 2023), indicating that behavioral impairments caused by developmental BaP exposure elicit multigenerational effects. These observations fuel the effort to resolve the mechanisms that underlie BaP neurotoxicity, which currently remains poorly understood, with the long-term goal of preventing or treating the neurotoxic effects of this ubiquitous pollutant. Recent work points to two possible variables that may play a role in defining how developmental BaP exposure ultimately gives rise to neurodegenerative disorders later in life: the gut microbiome and the aryl hydrocarbon receptor (AHR).

The gut microbiome may be an important component of the host’s biology that interacts with various chemical exposures, and researchers are just beginning to consider this component as part of the host’s response to toxicants (Bertotto, Catron, and Tal 2020; Claus, Guillou, and Ellero-Simatos 2016; Sharpton et al. 2021; Sutherland et al. 2020; Tu et al. 2020; Dong and Perdew 2020; Catron, Gaballah, and Tal 2019; Weitekamp et al. 2019; Chi et al. 2021; Maurice, Haiser, and Turnbaugh 2013; Stagaman, Sharpton, and Guillemin 2020). The gut microbiome coordinates with the central nervous system through a reticulated axis of communication that is collectively referred to as the gut-brain axis (Ahmed et al. 2022; Appleton 2018; Bertotto, Catron, and Tal 2020; Carabotti et al. 2015; Cryan et al. 2019; Dempsey, Little, and Cui 2019). The establishment of dysbiotic gut microbial communities has been shown to drive neurodegenerative phenotypes in mice and humans (Toledo et al. 2022; Singh, Dawson, and Kulkarni 2021; Korf, Ganesh, and McCullough 2022) due to perturbations to the homeostatic contribution of the gut microbiome to neurophysiology. Early-life exposures to environmental toxicants are theorized to perturb gut microbiome assembly and drive the successional development of dysbiotic communities. Indeed, our prior work found that early-life exposure to BaP perturbed gut microbiome assembly in zebrafish larvae (Stagaman et al. 2024). However, it remains unknown if these effects ultimately manifest as differential microbial communities among developed (adult) fish, and if such effects persist in subsequent generations.

While the AHR pathway is a known mechanism underlying BaP genotoxicity, its role in BaP neurotoxicity remains poorly defined and of interest (Chepelev et al. 2015). Recent research using AHR mutant zebrafish suggests that the receptor plays a role in larval hyperactivity resulting from early-life BaP exposure (Knecht et al. 2017a). The AHR binds BaP directly as part of the regular host-toxicant response, and in doing so, initiates host metabolic pathways that biotransform BaP into catabolites that are themselves neurotoxic (Bukowska, Mokra, and Michałowicz 2022; Saunders et al. 2006; Chepelev et al. 2015). Moreover, the AHR is a mechanism through which hosts and their gut microbiota interact: AHR binds gut microbial metabolites like tryptophan, inducing subsequent shifts in gut microbiome composition (Dong and Perdew 2020; Dong et al. 2020; L. Zhang, Nichols, and Patterson 2017; L. Zhang et al. 2015; Ma et al. 2020; Shankar et al. 2020; Sun et al. 2020). These observations raise the possibility that AHR may be a critical determinant of how BaP elicits neurotoxicity and, further, may interact with the gut microbiome in ways that affect this neurotoxic response. To our knowledge, no work to date has considered the effect of AHR on the gut microbiome and neurotoxicity in the context of developmental BaP exposure.

In this study, we sought to evaluate whether the AHR pathway and gut microbiome are linked to BaP neurotoxicity as well as its generational effects. To do so, we leveraged the zebrafish model, which affords a tractable and scalable means of controlling embryonic exposure, and specifically evaluated the gut microbiome and behavior of fish (F0) developmentally exposed to BaP, as well as their offspring (F1) and a subsequent generation (F2) of these fish. Because zebrafish grow rapidly, they are amenable to studies that investigate the developmental and generational consequences of embryonic exposure. Additionally, morpholino technologies have been developed that enable manipulation of specific zebrafish genes precisely during only the exposure period. Here, we utilized a morpholino of the AHR2 gene, which is the zebrafish homologue responsible for binding BaP, to transiently knock down embryonic expression during development in F0 generation fish. This manipulation was concurrent with exposure to 10 uM of BaP, which our prior work found was sufficient to elicit neurodevelopmental and generational effects in zebrafish (Knecht et al. 2017b). Specifically, we exposed developing zebrafish embryos to either 10 uM BaP (AHR2Mo-/BaP+), a morpholino that transiently blocks developmental AHR2 expression (AHR2Mo+/BaP-), both the toxicant and morpholino (AHR2Mo+/BaP+), or relative controls (AHR2Mo-/BaP-) from 0 to 5 days post fertilization (Fig 1). The four treatments were then raised in chemical-free water to assess potential developmental effects (in adult F0 fish). We also evaluated effects in adult F1 and F2 fish to test for potential multigenerational effects, where the F1 generation represents an intergenerational mechanism and F2 represents a transgenerational mechanism in zebrafish. Our analysis uncovered a link between BaP exposure, microbiome composition and functional capacity, and zebrafish behavior, and found that AHR2 was a key mediator of these associations. Our work also produced evidence that indicates that the effect of developmental BaP or AHR2 morpholino exposure affects the microbiome and behavior of subsequent generations. Ultimately, our study supports the hypothesis that the gut microbiome and AHR pathways are components underlying BaP’s mechanisms of developmental and generational neurotoxicity.

**Figure 1:**
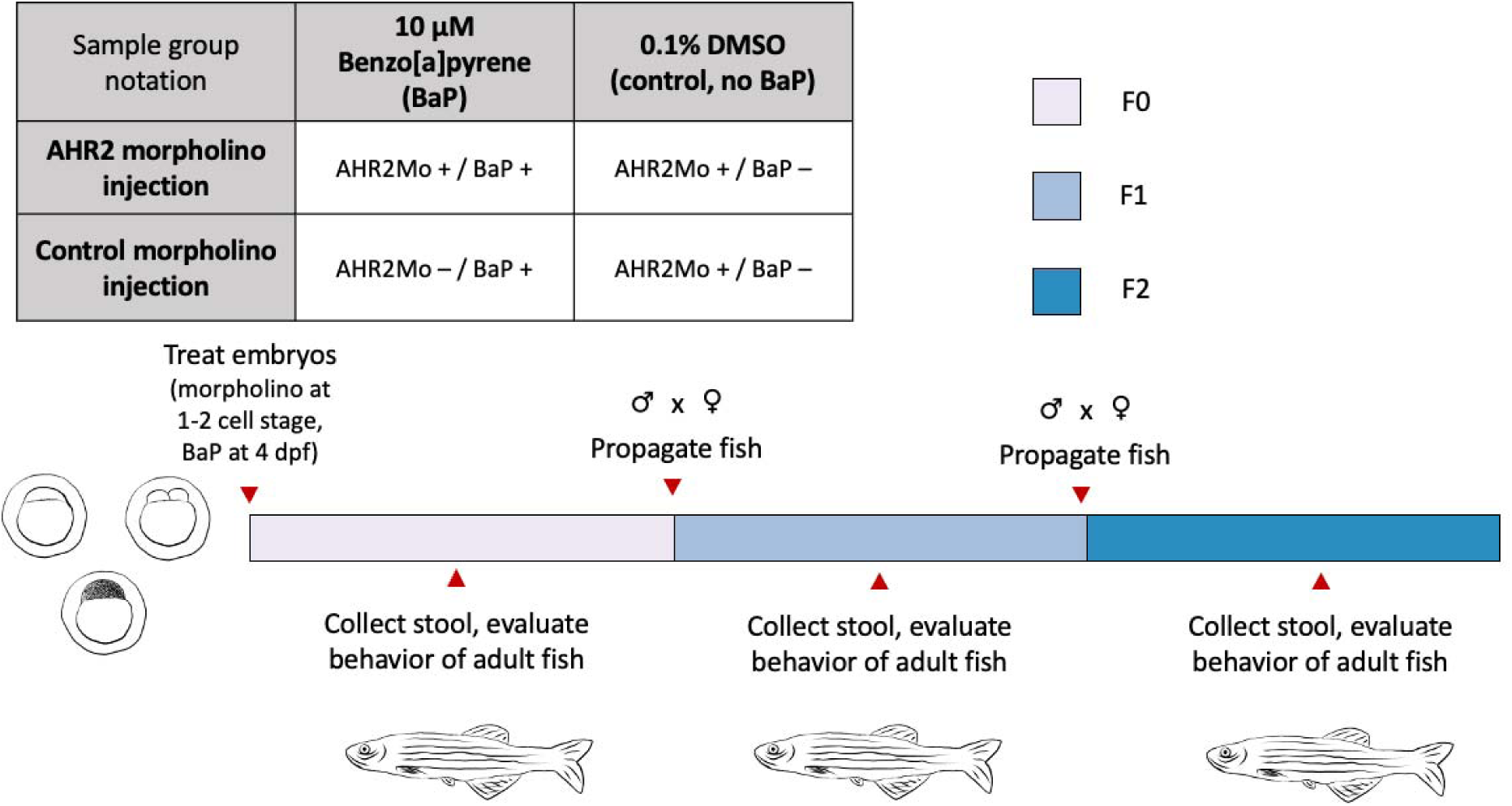
The factorial experimental design and workflow used in this study. The table at the top left shows the sample groups representing treatments we applied to zebrafish embryos in the F0 generation. The notation in the table is what is used in graphs and in text throughout the paper. The bar represents the workflow, with colors showing each generation and red arrows with text for relevant procedures and collections when they occurred.

## Results

### Overview

To investigate the role AHR and the gut microbiome, as well as their interaction, play in BaP neurotoxicity, embryonic zebrafish had their AHR2 gene transiently knocked down using a morpholino injection (AHR2Mo+) or were alternatively injected with a control morpholino (AHR2Mo-) (Fig 1). At 4 hpf, fish from these populations were either exposed to 0 uM BaP (AHR2Mo+/BaP- and AHR2Mo-/BaP-) or 10 uM of BaP (AHR2Mo+/BaP+ and AHR2Mo-/BaP+). Embryos were statically exposed to BaP in individual wells of a 96-well plate until 120 hpf, at which point the larvae were transferred to chemical-free water and raised until adulthood (∼4-5 months; F0) before undergoing adult behavioral assessments and gut microbiome analysis. Their offspring (F1) were reared and developed into adulthood in chemical-free water and underwent the same assessments as F0. From these F1 fish, F2 progeny were reared, developed into adulthood in chemical-free water, and subject to the same assessments.

### Transient suppression of AHR2 during development results in anxiety and altered social behavior

Previous studies assessing the impacts of developmental exposure of BaP on anxiety and social behaviors in adult zebrafish showed that the changes in behavior were heritable (Knecht et al. 2017b). In this study, we evaluated social perception using adult shoaling behavior metrics – inter-individual distance (iid) (cm), nearest neighbor distance (nnd) (cm) and shoaling speed (cm/min) – and evaluated anxiety using the free swim distance (cm) of adult fish when introduced into a new tank (Figs 2A, 3, and S1; Table S1). Taken together, perturbations in any of these metrics relative to the control group (AHR2Mo-/BaP-) zebrafish would indicate that behavior is altered.

**Figure 2:**
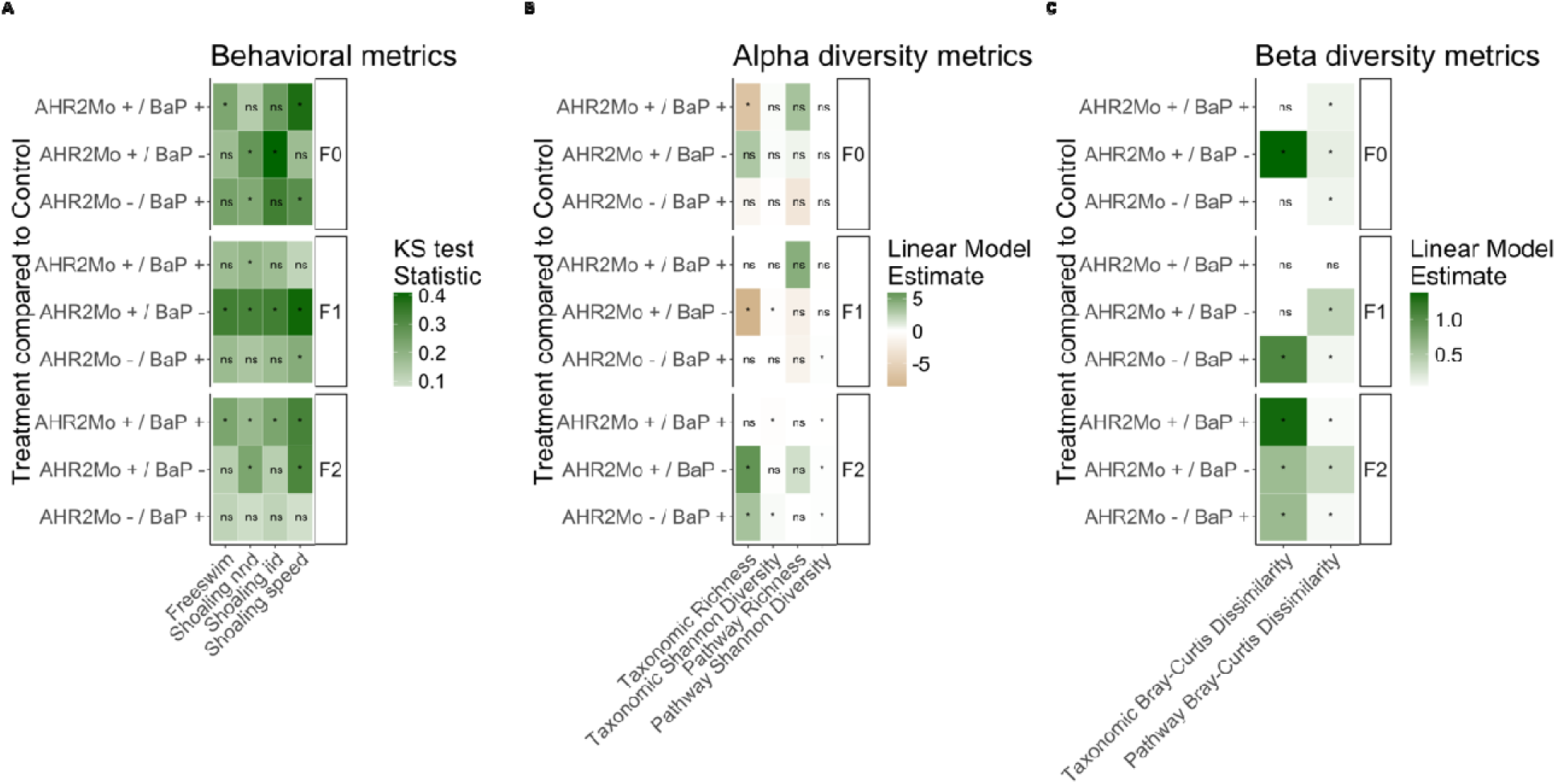
Summary heat map of statistical outcomes from behavioral and gut microbiome analyses. (A) shows the behavioral metrics and how they associate with each treatment (compared to the control, AHR2Mo-/BaP-), per generation. The KS statistic was used and is shown in the color gradient. (B) shows gut microbiome alpha diversity metrics (based on 16S [taxonomic] and shotgun metagenomic [pathway] data) associated with the treatments, per generation. (C) shows the gut microbiome beta diversity metrics (again, for 16S and shotgun metagenomic data) associated with the treatments, per generation. In Panels B and C, the color gradient represents the linear model estimate. Tables S2 and S3 (rows labeled “F0”) show expanded model results.

**Figure 3:**
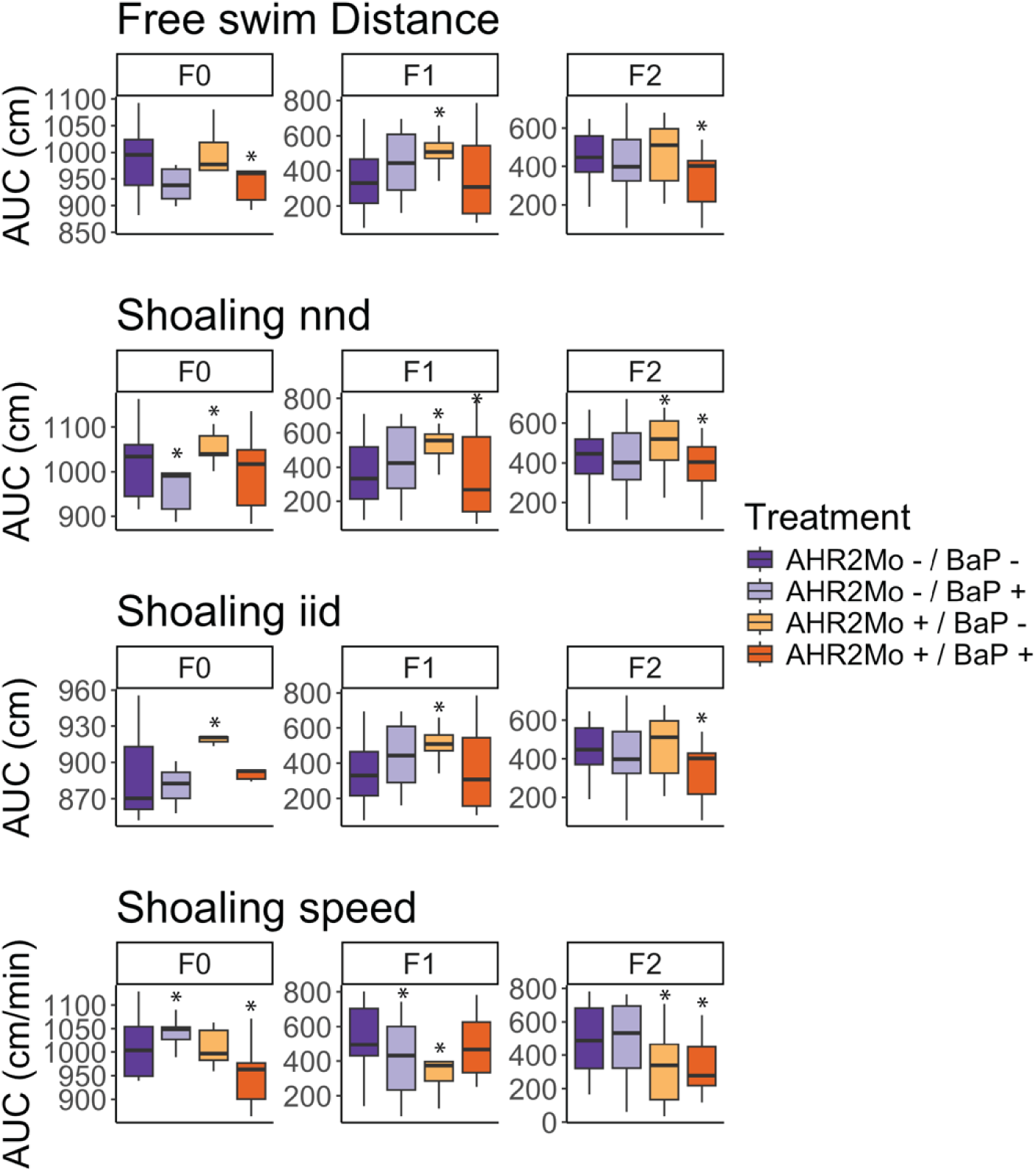
Differences in zebrafish behavior across experimental groups and generations. Four behavior metrics (free swim distance, shoaling nnd, shoaling iid, and shoaling speed) were measured using a ZebraBox camera, then used to create a cumulative density function (CDF) distribution for each treatment, per generation. CDF distributions were evaluated via a Kolmogorov-Smirnov (KS) test comparing each treatment CDF distribution to that of the control (AHR2Mo-/BaP-), for each generation, per metric. Here, we present box and whisker plots of the area under the curve (AUC) of the CDF distribution for each treatment, organized by generation and metric, to portray trends from the KS test visually. The asterisks indicate significance (p ≤ 0.05) in treatments relative to the control (AHR2Mo-/BaP-, dark purple), derived from the KS test described above (Table S1). Each color indicates the treatment. The center line of each box represents the median and outliers are not shown. Note that the y-axes for each plot show different scales, in order to more easily visualize trends within each data subset.

F0 fish showed several disturbed behavior metrics due to AHR2Mo+ treatment. We found that the 2 treatment groups not exposed to BaP (AHR2Mo-/BaP- and AHR2Mo+/BaP-) exhibited statistically different social behaviors based on AHR2Mo variation. The control group had a wider spread of shoaling cohesion than the fish with a transient knockdown of AHR2 during development (Figs 2A and 3, Table S1). These AHR2Mo+ fish exhibited increased median iid (KS test: p = 0.02) and nnd (KS test: p = 0.004), which indicates a less cohesive shoal, and this trend was persistent in F1 and F2 fish (Figs 2A and 3, Table S1). F1 and F2 fish also exhibited a slower shoaling speed (KS test: p = 0.01 and 0.00001, respectively) (Figs 2A and 3, Table S1). Hypoactive swimming and the desire to be in a less cohesive shoal indicates that the AHR2Mo treatment is associated with anxiety-like behavior that was heritable across generations.

When considering BaP’s impact on neurobehavior across a generation independently of AHR2 (AHR2Mo-/BaP+), the results recapitulated our previous study that demonstrated a change in social perception in F0 (KS test: decreased nnd; p = 0.03; higher shoaling speed; p = 0.02) (Figs 2A and 3, Table S1). The F1 generation did not show as robust a social perception change as F0, with the only statistically significant changes observed relating to a decrease in shoaling speed (KS test: p = 0.02) (Figs 2A and 3, Table S1). However, by F2, the abnormal social behavior was not detected. Since this behavior was not measured in our previous study, there was no way to compare the concordance of this result.

BaP and AHR2 morpholino treatment perturbed movement and social cohesiveness in zebrafish, and these effects manifested over generations. The role of AHR2 in BaP neurotoxicity was evaluated by comparing the AHR2Mo+/BaP+ treatment group and the control (AHR2Mo-/BaP-). There was an impact on the F0 social cohesion (KS test: slower shoaling speed; p = 0.00004) and hypoactive swimming (KS test: p = 0.05) (Figs 2A and 3, Table S1). F1 and F2 exhibited disrupted social behaviors and swimming behaviors as well. F1 fish exhibited a slightly lower median nnd with more variation (KS test: p = 0.05) but no change in swimming behavior (Figs 2A and 3, Table S1). By F2, the fish exhibited hypoactive swimming and decreased social cohesion (KS test: free swim, p = 0.004; nnd, p = 0.05; iid, p = 0.004; speed, p = 0.000009) (Figs 2A and 3, Table S1). Collectively, F0 and F2 both exhibited hypoactive swimming and decreased social cohesion, which are markers of anxiety-like behavior.

### Developmental BaP exposure, AHR2 disruption, and their interaction affect gut microbiome metrics in F0 fish

To determine if the microbiome is linked to BaP and AHR2 morpholino exposure, we first evaluate how the microbiome differs across experimental groups in fish directly exposed to these conditions (i.e., F0 fish). In particular, we developed statistical regression models that linked measures of microbiome biodiversity and composition to BaP exposure, AHR2 morpholino, sex, or the interactions between each pair of these. We calculated these metrics for two datasets: ASVs and metabolic pathways encoded in the metagenome, derived from 16S and shotgun metagenomic sequence data, respectively. Across these analyses, several overarching patterns emerged in the data.

First, we found that BaP exposure elicits modest effects on the gut microbiome. Neither taxonomic alpha-diversity nor beta-diversity was associated with BaP exposure as a main effect (Figs 2B and 2C). However, BaP exposure was associated with bacterial metabolic pathway beta-diversity (PERMANOVA: R^2^ = 0.06, p = 0.001), and metabolic pathway richness approached significance (linear repression: coefficient = - 2.8, p = 0.09). Beta dispersion was not associated with any experimental variables in taxonomic or pathway gut microbiome datasets (Tables S2 and S3).

That said, BaP elicits stronger effects on the gut microbiome in an AHR2-dependent manner. For example, the interaction between AHR2 morpholino treatment and BaP exposure was correlated with ASV richness (linear regression: coefficient = - 6.8, p = 0.002), whereas each individual variable (AHR2 morpholino and BaP exposure) was not significant (Fig 2B, Table S2). The control cohort (AHR2Mo-/BaP-) had the highest ASV richness, whereas AHR2Mo+/BaP+ fish had the lowest (Fig 4A). In addition, the interaction between BaP exposure and AHR2 morpholino was also associated with bacterial pathway beta-diversity (PERMANOVA: R^2^ = 0.07, p = 0.001) (Figs 2C, 4C, and S2B, Table S3). There was no effect of the interaction on beta dispersion (Tables S2 and S3). Taken together, these results point to an AHR2-dependent BaP exposure effect on the gut microbiome.

**Figure 4:**
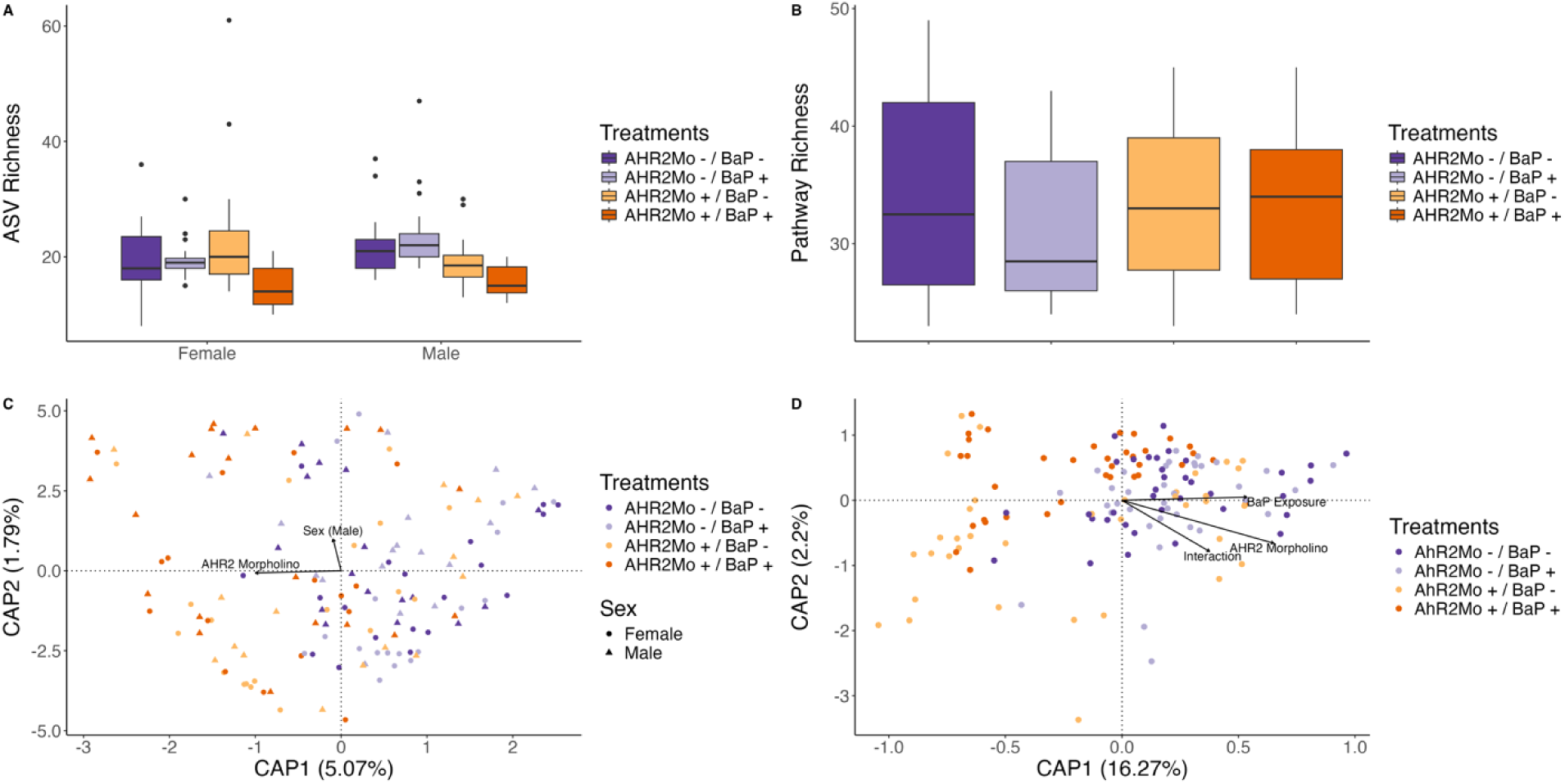
Plots showing alpha and beta diversity metrics in taxonomic and pathway data per significant treatments (identified via regression approaches) for generation F0. (A) is a box and whisker plot showing the ASV richness for each treatment (color) per sex (linear regression: exposure and morpholino interaction, coefficient = -6.8, p = 0.002; morpholino and sex interaction, coefficient = -5.3, p = 8.3 × 10^-5^). (B) is a box and whisker plot showing pathway richness for each treatment (color) (linear regression: no significant variables). (C) is a dbRDA ordination showing taxonomic beta diversity (Bray Curtis dissimilarity) per treatment (color) and sex (shape) (PERMANOVA: p = 0.001 and 0.03, respectively). (D) is a dbRDA ordination showing the pathway beta diversity (Bray-Curtis dissimilarity) per treatment (color) (PERMANOVA: p = 0.001 for all three covariates on the vectors).

In addition to influencing the effect of BaP on the microbiome, we found that AHR2 morpholino treatment elicited an effect on the zebrafish gut microbiome independently of BaP exposure. In fact, the effect of AHR2 morpholino treatment on the gut microbiome was stronger in many cases than that of BaP exposure (Figs 2A, 2B, 4A, and 4B). ASV richness and Shannon diversity were each associated with morpholino treatment, however this association differed as a function of sex (linear regression: richness, coefficient = -5.3, p = 0.01; Shannon diversity, coefficient = -0.5, p = 8.3 × 10^-5^) (Figs 2B, 4A, and S2). For example, ASV Shannon diversity in female AHR2Mo+ fish was higher than that of female AHR2Mo-zebrafish, whereas this pattern was reversed in males (Figs 2B and S3). Moreover, ASV Bray-Curtis dissimilarity was associated with AHR2 morpholino treatment (PERMANOVA: R^2^ = 9.3, p = 0.001) (Figs 2C and 4B). Pathway Bray-Curtis dissimilarity was also associated with morpholino treatment (PERMANOVA: R^2^ = 16.2, p = 0.001) (Fig 2C), as was beta-dispersion, albeit more weakly (linear regression: coefficient = 0.01, p = 0.052) (Figs 3C and S2B).

Supporting the above findings, several ASVs and microbial gene pathways in the zebrafish gut were correlated with AHR2 morpholino status and the interaction between BaP exposure and AHR2 morpholino status (but not BaP exposure alone) (Fig 5, F0 labels in heatmap). One example is *Shewanella*, a member of the core zebrafish microbiome (Sharpton et al. 2021; Roeselers et al. 2011), whose relative abundance increased in association with developmental BaP exposure in an AHR2-dependent manner (Wald test: interaction between exposure and morpholino, p-value ≤ 0.05) (Fig 5A). Additionally, *Rheinheimera*, *Pseudoduganella* and *Chitinimonas* relative abundances decreased in association with morpholino and exposure treatment together (Wald test: p-value ≤ 0.05), while *Plesiomonas* relative abundance increased with morpholino treatment (Wald test: p-value ≤ 0.05) (Fig 5A). We also detected bacterial pathways related to xenobiotic and neurotransmitter (e.g., Gamma-aminobutyric acid [GABA]) metabolism, DNA methylation, fermentation, short chain fatty acid synthesis and degradation, and cell growth (TCA cycle and cell membrane metabolism especially) that are enriched or depleted in association with AHR2 morpholino treatment and its interaction with BaP exposure, but not exposure alone (Fig 5B, Table S4). These results further support the finding that specific ASVs and pathways in the gut microbiome respond to BaP exposure in an AHR2-dependent manner and AHR2 itself has a strong effect on gut microbiome succession from embryo to adulthood.

**Figure 5:**
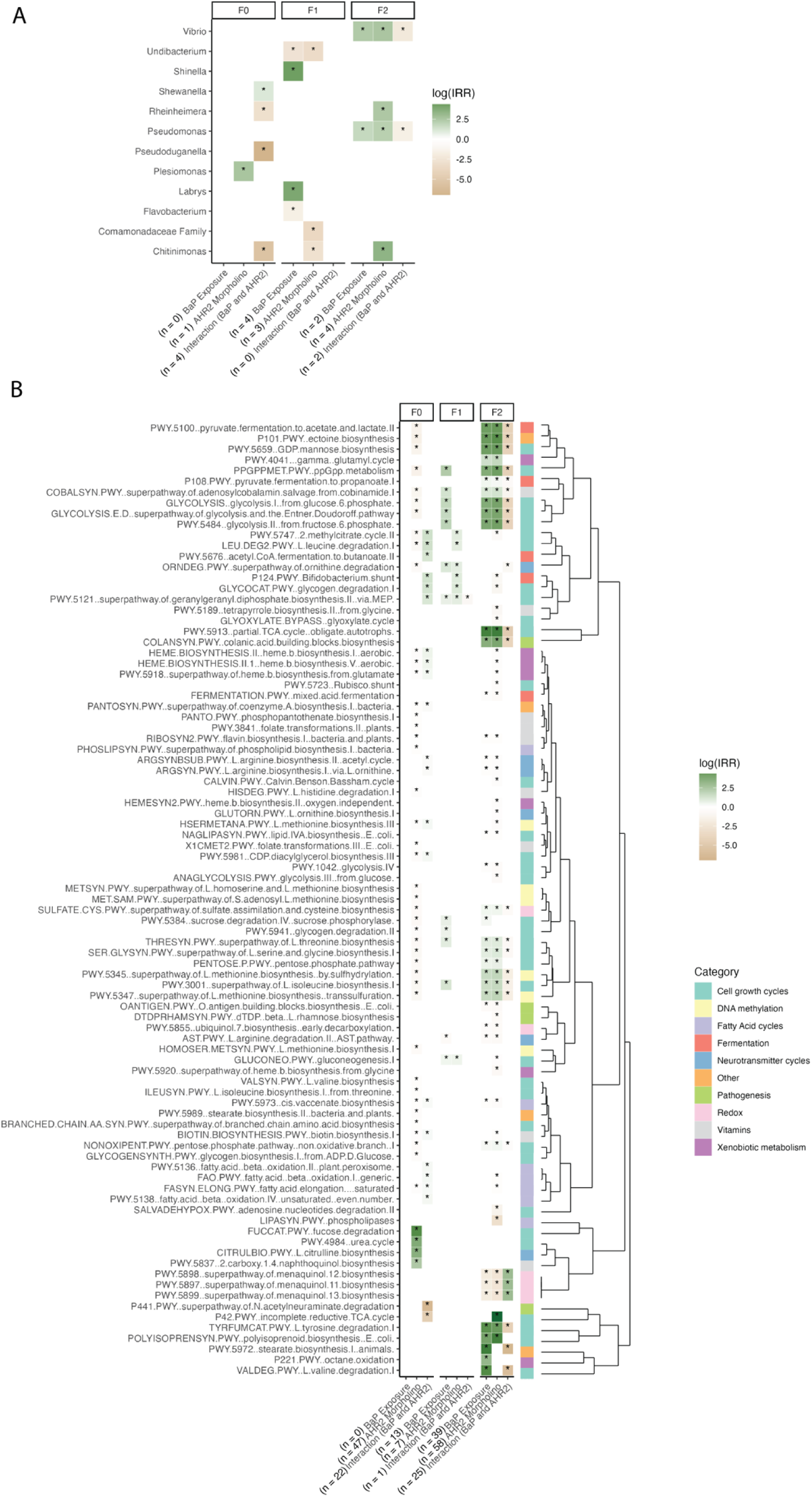
A heatmap of indicator taxa and pathways associated with model terms, per generation. (A) shows indicator taxa listed on the y-axis while (B) shows the indicator pathways, organized by a dendrogram based on the Bray-Curtis matrix (right), and each organized by generation (top labels). Each y-axis lists the model terms used in a negative binomial model of each taxon/pathway relative abundance, along with the number of significant associations shown in parentheses. Log(IRR) is represented by a color scale, which is the strength and direction of the association wherein positive numbers are dark green and a positive association between the taxon/pathway relative abundance and the model term, and vice versa for negative numbers. The bottom set of colors in panel B indicates the broad category each pathway falls under (curated using available information from BioCyc and literature searches). Asterisks represent significant p-values from the model.

### Gut microbiome and behavior changes associated with BaP exposure and AHR2 morpholino persist over subsequent generations

We found that, overall, patterns of association between experimental factors and gut microbiome metrics persisted in subsequent generations. Specifically, we found that BaP exposure, AHR2 morpholino status, and the interaction between the two had a strong association with alpha- and beta-diversity metrics in subsequent generations. In some cases, these effects were stronger than was observed in F0 fish. We used two modeling approaches with this data: (1) the same modeling approach as in F0 fish applied to each subsequent generation (F1 and F2), wherein BaP exposure, AHR2 morpholino, sex, and the interaction between each of these were independent variables and alpha- and beta-diversity gut microbiome metrics were each the dependent variable, and (2) using all the data (i.e., with all three generations present) and included “generation” as an independent variable in all models, along with its interaction with BaP and AHR2 morpholino treatments. These approaches were each applied to 16S and metagenomic datasets. The results of both modelling approaches were congruent with each other. For simplicity, we present the results of the first approach in the main text. Tables S2 and S3 show all model results from both approaches and all gut microbiome metrics for reference. In the following paragraphs, we expand on the specifics of these patterns.

Linear regression models showed that BaP exposure and AHR2 morpholino treatments in F0 embryos are overall strongly associated with the alpha-diversity of gut microbiomes in F1 and F2 fish. In F1 fish, ASV richness and Shannon diversity were both associated with AHR2 morpholino treatment (richness coefficient = -8.6, Shannon diversity coefficient = -0.2, p = 0.005 for both metrics), but not exposure (Figs 2B and 6A; Table S2). By contrast, metabolic pathway Shannon diversity was associated only with BaP exposure (coefficient = 0.07, p = 0.0005) (Figs 2B and 6B; Table S2). In F2 fish, ASV alpha diversity was associated with morpholino status (richness, coefficient = 5.8, p = 1.2 × 10^-7^), exposure (richness coefficient = 3.2, Shannon diversity coefficient = 0.3, p = 0.002 for both), and their interaction (Shannon diversity, coefficient = -0.2, p = 0.05) (Figs 2B and 6A; Table S2). Pathway Shannon diversity of F2 fish microbiomes was associated with BaP exposure (coefficient = 0.06, p = 0.0002), morpholino status (coefficient = 0.06, p = 0.0002), and their interaction (coefficient = -0.06, p = 0.01) (Figs 2B and 6B; Table S2).

**Figure 6:**
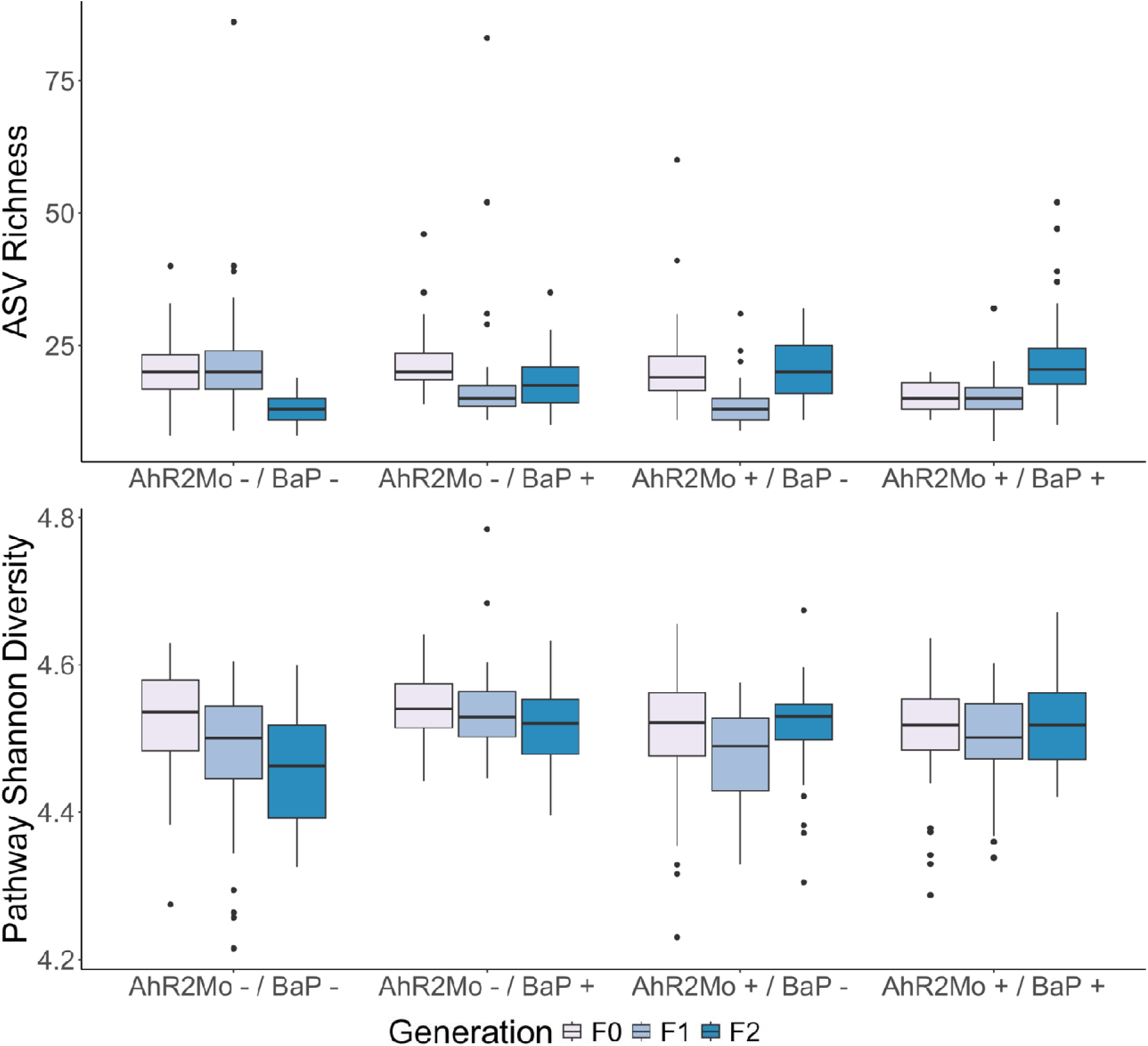
Box and whisker plots of alpha diversity metrics per treatment in each generation, for taxonomic and pathway diversity. (A) Taxonomic richness and (B) pathway Shannon index were the only significant metrics among all alpha diversity metrics tested (see Table S1 for statistics). Color in both graphs shows generations F0 – F2 and x-axis lists the different treatments.

PERMANOVA models of beta-diversity also support the finding that BaP and AHR2 perturbations have sustained effects on gut microbiome composition and functional capacity in subsequent generations. Pathway Bray-Curtis dissimilarity was associated with morpholino status (R^2^ = 0.17, p = 0.001) and BaP exposure (R^2^ = 0.03, p = 0.002) (Figs 2C, 7A, and S3B; Table S3). Further, we observed that all treatments deviated from the control in terms of median Bray-Curtis dissimilarity in F1 and F2 generations relative to F0 (Fig 7B; Table S3). F1 was particularly dissimilar from F0, whereas F2, despite being further removed from F0, was more similar to F0 than F1 was, congruent with a transgenerational effect of BaP and AHR2 on gut microbiome beta diversity (Fig 7B; Table S3). Beta dispersion was associated with BaP exposure and AHR2 morpholino treatment in the F1 generation only (linear regression; BaP coefficient = 0.03, p-value = 0.04; AHR2 coefficient = -0.03, p-value = 0.02) (Fig 7C; Table S3). ASV Bray-Curtis dissimilarity in F1 fish was associated with BaP exposure (R^2^ = 0.04, p = 0.001) (Fig S3A; Table S3). In F2 fish, pathway Bray-Curtis dissimilarity in F2 fish was also associated with morpholino status (R^2^ = 0.21, p = 0.001), BaP exposure (R^2^ = 0.03, p = 0.002), and their interaction (R^2^ = 0.02, p = 0.004) (Fig S3B), and beta-dispersion of pathways was associated with morpholino status (linear regression: coefficient = 0.02, p = 0.01) and BaP exposure (linear regression: coefficient = 0.02, p = 0.02) (Table S3). ASV Bray-Curtis dissimilarity was associated with morpholino status (R^2^ = 0.06, p = 0.001), BaP exposure (R^2^ = 0.03, p = 0.005), and their interaction (R^2^ = 0.03, p = 0.02). Beta-dispersion of ASVs in F2 fish were associated with morpholino status (linear regression: coefficient = 0.09, p = 0.005) (Table S3).

**Figure 7:**
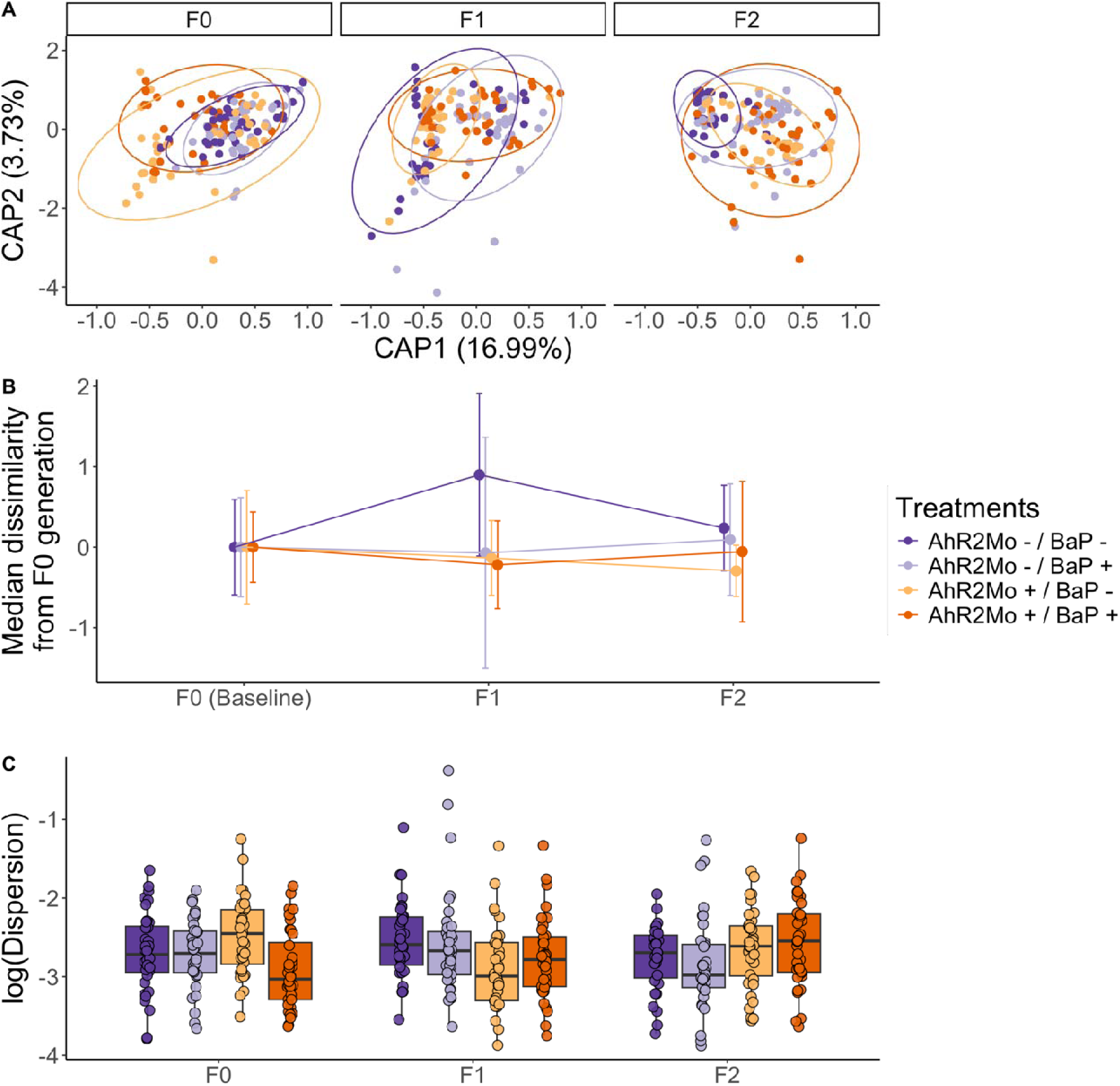
Bray-Curtis dissimilarity of functional pathways across generations and treatments across the whole dataset. (A) represents a dbRDA ordination where each point is a zebrafish gut microbiome sample, with color showing treatments and the ordination is faceted by generation. The ellipses show each generation per treatment grouping (n = 12) where color is treatment. (B) is a dot plot of the median distance from the points in each generation-treatment combination to the F0 centroid for the corresponding treatment (see methods for details). Distances are taken from the dbRDA ordination in Panel A, normalized to the F0 centroid, and parsed by generation and treatment (color). Lines colored by treatment connect each treatment across generations. (C) is a box and whisker plot of the dispersion values for each treatment (colors) within each generation. Dispersion values have been log transformed for ease of visualization. The legend is common to all three panels.

Several ASVs and metabolic pathways were associated with BaP exposure, AHR2 morpholino status, and their interaction within each generation (Fig 5). For example, the Comamonadaceae family, a core zebrafish group (Sharpton et al. 2021), exhibited decreased relative abundance with AHR2 morpholino treatment in generation F1 (Wald test: p-value ≤ 0.05) (Fig 5A). In the F2 generation, ASVs from the core zebrafish gut genera *Vibrio* and *Pseudomonas* (Sharpton et al. 2021) both exhibited increased relative abundance when exposed to BaP and the AHR2 morpholino (Wald test: p-value ≤ 0.05), and had decreased relative abundance in association with treatment with both BaP and AHR2 morpholino (p-value ≤ 0.05) (Fig 5A). Further, there were several pathways that were associated with BaP and/or AHR2 morpholino to various extents (Fig 5B), again including pathways linked to xenobiotic metabolism, neurotransmitter metabolism, DNA methylation, fermentation, short chain fatty acid metabolism, and cell growth. Extended explanations of each pathway as well as the superclasses from BioCyc can be found in Table S3. Broadly, the associations were stronger and increased in the F2 generation (and weak or not at all present in the F1 generation by comparison, and less so in F0) or were maintained between subsequent generations, as is shown in the enumerated differentially abundant pathways listed in the x-axis of Figure 5B. This observation is consistent with the alpha-and beta-diversity results pointing to intergenerational effects of BaP exposure and AHR2 perturbation on the zebrafish gut microbiota, spanning differences in taxonomic composition as well as functional capacity (Fig 5B).

## Discussion

Our investigation sought to address three key questions: (1) how exposure to BaP and developmental expression of AHR2, a key gene in the zebrafish AHR pathway, links to adult zebrafish behavior, (2) how these same variables associate with the structure and function of the adult zebrafish gut metagenome, and (3) whether these associations subsequently manifest across three generations of fish. We found that exposure to BaP had a modest effect on zebrafish behavior and gut microbiome, but this effect was stronger when the AHR2 gene was perturbed, implying that BaP affects zebrafish behavior and the gut microbiome (composition, structure, and functional capacity) in an AHR2-dependent manner. These associations also persisted and, in some cases, strengthened over subsequent generations (F1 and/or F2). Lastly, we found that AHR2 had a strong effect on zebrafish behavior and gut microbiome assembly independently of BaP exposure across all generations. Overall, these results support the hypothesis that BaP perturbs the successional development the gut microbiome to dysregulate behavior in an AHR2-dependent manner. Below, we expand on the major patterns we observed in the data and their implications, given prior research.

Embryonic BaP exposure elicits modest effects on zebrafish gut microbiome and behavior in adult zebrafish. In our analysis of fish directly exposed to BaP (i.e., F0 AHR2Mo-zebrafish), we found that BaP elicits significant effects on two measures of adult fish behavior: shoaling nnd and shoaling speed (Fig 2A). The most directly comparable past study we know of uncovered a stronger BaP effect on specific behavior measures, like shoaling nnd and iid, but ultimately found the same outcome – that social behavior is perturbed by BaP (Knecht et al. 2017b). This slight difference could also stem from updated methodology in the behavior assays. Beta diversity of the gut microbiome’s functional capacity in F0 fish shifted with BaP exposure as well (Fig 2C). This result indicates that, like behavior, the successional dynamics of the gut microbiome over the course of fish development are modestly impacted by embryonic BaP exposure. Past research also measured a modest effect of embryonic BaP exposure at increasing doses on larval zebrafish gut microbiome structure and composition (Stagaman et al. 2024). Another study found similarly modest shifts in gut microbiota and moderate inflammation of the gut epithelium in mice that were dosed with BaP in adulthood and then promptly measured (Ribière et al. 2016). However, this study differed from ours in that they didn’t measure the effect of developmental BaP exposure on the adult gut microbiome or behavior.

In contrast to the modest effects described above, exposing embryos to both BaP and the AHR2 morpholino results in a strong effect on both behavior and the gut microbiome. As far as we are aware, no study has previously tested the effect of early life AHR perturbation alongside developmental BaP exposure on the gut microbiome. In many instances, we found that the effect of BaP on the gut microbiome is sensitive to the state of AHR2 functioning at the time of exposure (Fig 2). For instance, a large number of bacterial pathways involved in xenobiotic response and gut-brain crosstalk were not differentially abundant in association with BaP exposure unless AHR2 was also disrupted (Fig 5B). This result indicates that the AHR2 expression state at the time of BaP exposure dictates how the host and its microbiome respond to BaP exposure. The importance of AHR2 functioning makes sense because it binds numerous environmental chemicals, not only BaP, in addition to microbially derived metabolites (Antkiewicz, Peterson, and Heideman 2006; Brown et al. 2015; Knecht et al. 2017a; Claus, Guillou, and Ellero-Simatos 2016; Goodale et al. 2015; Dong et al. 2020; Ma et al. 2020). Additionally, our prior work found that developmental BaP exposure elicits a modest effect on the gut microbiome assembly in larval zebrafish (Stagaman et al. 2024). BaP binds to AHR2, resulting in DNA adducts that are sometimes not cleared and themselves are related to neurotoxicity (Perera et al. 2012), so it is possible that the reason we see an interaction effect of BaP and AHR2 on the microbiome is attributable to the variation in these BaP catabolites, which may perturb the microbial community.

Our study also uncovered evidence that developmental BaP exposure can elicit multigenerational effects on the gut microbiome and behavior, and that these effects are mediated alongside developmental AHR2 expression state. For example, BaP exposure exhibits stronger effects on behavior and gut microbiome metrics in F1 and F2 fish compared to F0 fish. Specifically, BaP exhibits an inter- and transgenerational effect on gut microbiome alpha- and beta-diversity metrics, many taxonomic and pathway bacterial relative abundances, and a variety of behavior measures. These multi- generational effects were even larger in fish subject to AHR2Mo+ injection, indicating that AHR2 may influence how BaP manifests its effects across generations.

This is the first study to demonstrate that early life exposure to BaP has a generational behavior effect that is dependent on AHR2 in zebrafish and that these effects are accompanied by perturbations to the gut microbiome. Generational effects such as those observed here imply an epigenetic mechanism may be at play. We found that bacterial pathways related to methylation cycles were differentially abundant in association with BaP and/or AHR2 perturbation, within each generation (Fig 5). Past research also found that BaP metabolism by AHR leads to the production of DNA adducts and alterations to the DNA methylome (Knecht et al. 2017b; Gao et al. 2018; Wilson 2023; Fang et al. 2013). BaP also induces multigenerational behavioral effects in association with changes in zebrafish genome-wide DNA methylation (Pandelides et al. 2023; Knecht et al. 2017a). Notably, both gut microbiome and behavior manifested transgenerational effects of AHR2 perturbation, supporting the idea that epigenetics is at play to some extent. Future work should clarify the role of exposure-induced variation on the epigenome of zebrafish and the gut microbiome, and the consequences on the microbiome, host behavior, and the gut-brain axis.

Additionally, developmental AHR2 state is associated with the behavior and gut microbiome in adult zebrafish and their progeny. We found that developmental AHR2 morpholino treatment perturbs adult behavior and gut microbiome assembly, sometimes more strongly than BaP exposure, and this effect spanned all three generations (Fig 2). AHR regulates gut homeostasis by binding bacterially produced metabolites of tryptophan, mainly indoles, kynurenine, and serotonin (Dong et al. 2020; Dong and Perdew 2020; Ma et al. 2020), and AHR2 function in zebrafish is required for normal behavioral development (Garcia et al. 2018). AHR ligands like tryptophan and indole compounds have also been associated with neurodegenerative and neurological disorders and central nervous system inflammation (Dong and Perdew 2020; Ma et al. 2020; Rothhammer et al. 2016). Consistent with this previous research, we identified various bacterially derived neuromodulator pathways that were differentially abundant in association with AHR2 morpholino injection, including tryptophan, short-chain fatty acids, and GABA (a serotonin precursor) (Dong and Perdew 2020; Ma et al. 2020) (Fig 5B). One specific example is the L-Ornithine biosynthesis pathway (Table S3), which has slightly decreased abundance in F2 fish derived from F0 fish that had AHR2 morpholino treatment (Fig 5B). L-Orthinine biosynthesis increases the level of AHR ligand L-kynurenine, produced from tryptophan metabolism in gut epithelial cells (Qi et al. 2019). Additionally, one review hypothesized that BaP binds to AHR to then transcribe a N-methyl-d-aspartate glutamate receptor (NMDAR), which it turn mediates neurotoxic effects (Chepelev et al. 2015). To do this, NMDAR binds glutamate, and we identified differentially abundant pathways related to glutamate metabolism cycles in association with AHR2 (Fig 5B). These various lines of evidence together suggest that perturbing AHR2 function for a temporary moment in early life has some effect on the functional capacity of gut microbiota. These metabolites are not only important for behavioral development and establishing homeostasis between the gut and brain, but also aid in further assembly of the adult gut microbiome. The results underscore the importance of AHR2 to zebrafish development, particularly in the context of the gut-brain axis, and future work is needed to clarify the mechanisms through which AHR2 drives neurodevelopment.

Taken together, observations stemming from this study support the hypothesis that BaP exposure interacts with AHR2 to elicit neurotoxicity and dysregulate gut microbiome assembly and the gut-brain interaction. Our work also raises the hypothesis that environmental chemical exposure may elicit multi-generational dysbioses in the gut microbiome. Future work should characterize the mechanisms behind how different gut microbiota interact with BaP in the gut, whether BaP and microbial ligands somehow interact to bind to AHR2 in the context of zebrafish neurotoxicity, and how these effects propagate to subsequent generations. Future work should also consider other neurotoxic PAHs that bind AHR2, like TCDD, to determine if the interactions identified here manifest in other chemicals. Finally, our study highlights the utility of the zebrafish model in these efforts, as it offers tools to explore how host genetics interact with the gut microbiome to clarify neurotoxicity mechanisms, scalability to resolve otherwise subtle – but biologically important – effects, and rapid growth to afford insight into multi-generational effects.

## Methods

### Data collection and zebrafish husbandry

This experiment used the 5D zebrafish line, which is a genetically diverse line (Kent et al. 2011). Zebrafish were housed and maintained at Oregon State University’s Sinnhuber Aquatic Research Laboratory (SARL) using a published husbandry protocol (IACUC-2021-0166) (Kent et al. 2011; Barton, Johnson, and Tanguay 2016). Briefly, tanks were filled with high-quality groundwater, which was aerated, filtered for particulate matter, and UV sterilized before use. Recirculated water was filtered via reverse osmosis and activated carbon, and it was supplemented with Instant Ocean salts (Spectrum Brands, Blacksburg, VA, USA) and sodium bicarbonate as needed to maintain a pH of 7.4. Temperature was maintained at 28 ± 1 °C. For spawning the F1 and F2 generations, adult fish spawned in sex-gated tanks and embryos were collected immediately afterwards.

Morpholino and BaP exposures were performed on F0 generation embryos. We used an A-translation blocking morpholino (5′ TGTACCGATACCCGCCGACATGGTT 3′) and a standard control morpholino (5’ CCTCTTACCTCAGTTACAATTTATA 3’) purchased from Gene Tools (Philomath, OR, USA). One of these two morpholinos (∼2 nL of constituted morpholino sequence) was injected into the yolk of 1-2 cell stage embryos. Phenol red dye was used to confirm successful injection. Embryos were kept in E2 embryo medium (EM) consisting of 15 mM NaCl, 0.5 mM KCl, 1 mM CaCl2, 1 mM MgSO4, 0.15 mM KH2PO4, 0.05 mM Na2HPO4, and 0.7 mM NaHCO3 buffered with 1 M NaOH to pH 7.2 (Westerfield 2007). Embryos were then stored at 28 ± 1 °C in the dark until dechorionation and subsequent BaP exposure.

Embryos were dechorionated at 4 hours post-fertilization (hpf) for BaP treatment, using a custom pronase-based dechorionation instrument (Mandrell et al. 2012). Benzo[a]pyrene 10 mM stocks were made with 100% dry dimethylsulfoxide (DMSO) (Sigma-Aldrich, 96% purity). This BaP stock was water-bath sonicated for 20 minutes to ensure proper dissolution, then a working stock was made using EM and sonicated for 30 minutes. 100 µL of BaP and 0.1% DMSO control stocks were pre-loaded into the wells of round-bottom 96-well Falcon plates at concentrations of 10 µM BaP (normalized to 0.1% DMSO) or a 0.1% DMSO vehicle control. Embryos, either microinjected with either control or AHR2 morpholino, were then plated into individual wells of the plate and then moved out at 120 hpf to chemical-free EM to grow into adulthood. In adulthood, zebrafish gut microbiome and behavioral data was collected, as well as subsequent spawning and sampling of F1 and F2 generations.

Adult zebrafish were screened for standard behavioral changes at each generation (F0, F1, F2) using previously defined assays (Knecht et al. 2017b; Reif et al. 2016). Adult zebrafish were put into tanks alone and a Point Gray Camera was used in conjunction to Viewpoint ZebraLab 3.3 (Lyon, Fance) software to record the distance they swam in one-minute increments over the course of 30 minutes. Of note, the ZebraLab software v3.3 was used to measure F0 fish and was upgraded to v5.18 for F1 and F2, and accounts for the difference in sensitivity of measurements that is apparent in the presented analysis. Therefore, our analysis, detailed in a later subsection, focuses on finding trends between treatments within each generation. We measured four assays: (1) free swim distance (cm), a measure of distance swum when a fish is introduced into a new tank, thus representing the exploration of a environment; (2) shoaling nearest-neighbor distance (nnd) (cm), a social behavior metric measuring the distance to nearest neighbor in a group of four fish, where one fish is tracked; (3) shoaling inter-individual distance (iid) (cm), a social behavior measure of the distance between pairs of fish in a group of four fish; (4) shoaling speed (cm/min), a social behavior metric of the speed of a focal fish in a group of four fish. Consistent with past published work, dead or morphologically deficient fish were removed from the analysis, and the first 10 and last 8 minutes were trimmed from the data (fish tend to behave variably or freeze at the start because they’re in a new environment and acclimate to the new tank by the end of the measurement period).

Zebrafish were sampled for gut microbiome analysis in adulthood (at least 120 days post fertilization) at each generation and treatment group. Fish were euthanized using MS-222 at 470 mg/L for 10 minutes as per the accepted IACUC protocol. Fish were dissected to collect whole large and small intestines, which were stored in sterile 2.0 mL centrifuge tubes in liquid nitrogen immediately after collection and stored at -80C until all samples were collected and ready for DNA extraction in bulk. During dissection, fish metadata were collected: sex, length, and width.

### Processing gut microbiome sequences

Samples were prepared for 16S rRNA and shotgun metagenomic sequencing. We extracted DNA using the DNeasy PowerSoil Pro DNA Extraction kits (Qiagen, Hilden, Germany). The V4 region of the 16S rRNA was PCR amplified with EMP protocol and primers (citation). Prior to sequencing, we checked product on a X% agarose gel and pooled samples, normalizing concentrations using Qubit readings (Thermo Fisher Scientific, Waltham, Massachusetts, USA). Sequencing of the 16S rRNA region and shotgun metagenomic sequencing was done through the Center for Quantitative Life Sciences at Oregon State University on the Illumina MiSeq high-throughput sequencer (San Diego, California, USA) for 16S sequencing and an Illumina HiSeq 2000 sequencer (San Diego, California, USA) for shotgun metagenomic sequencing. To obtain higher sequence resolution in the metagenomic shotgun sequence dataset, the same samples were sequenced across multiple lanes, and we concatenated files from the same sample after demultiplexing. All sequences are publicly available in an NCBI sequence repository project (PRJNA######).

16S sequence processing was done with a modified DaDa2 pipeline in R developed by our lab for gut microbiome analysis (https://github.com/kstagaman/sharpton-lab-dada2-pipeline version 0.3.4) implemented in R (version 2022.12.0) (Beghini et al. 2021). Reads were trimmed to 225bp forward and reverse 200bp. Reads with a quality score less than 30 were thrown out. We assigned amplicon sequence variants (ASVs) with the Silva 16s database training set (version 138.1) (Quast et al. 2013). A phylogenetic tree was built with the Mothur (version 1.47.0) workflow (Schloss et al. 2009) using the NAST algorithm from FastTree2 (Price, Dehal, and Arkin 2009). Before diversity-based analyses, we filtered mitochondria and chloroplast sequences and samples that contained extremely low or no counts, and 16S rRNA data was rarefied to 10,000 sequences.

To process shotgun metagenomic sequences, we used the Sharpton lab metaGTXprocessing pipeline (https://github.com/kstagaman/sharpton-lab-metaGTx.processing) which is an adapted version of the BioBakery workflow implemented in R (version 4.1.2 (2021-11-01)) (Beghini et al. 2021). Briefly, the workflow proceeds as such: kneaddata (v0.10.0) removes contaminant reads that match the host (via Bowtie2) and trims sequences (via Trimmomatic); microbial taxonomy is assigned via Metaphlan (version 3); gene families and pathways are inferred using the assigned taxonomy using Humann (v3.0.0.alpha.3) (Beghini et al. 2021). This information was summarized in three tables: taxa abundance per sample, gene family abundance per sample, and pathway abundance per sample. We used all three for exploratory and supplementary analyses but focus on the pathway abundances per sample in the analyses described in the main text. Before diversity-based analyses, we removed samples that contained extremely low or no counts compared to the rest of the samples using the raw sequence count per sample frequency distribution.

### Analysis and statistics

We determined whether embryonic BaP exposure, AHR2 morpholino status, or their interaction were associated with alpha-diversity, beta-diversity, or beta-dispersion of ASVs (derived from 16S marker gene sequencing) and/or putative pathways (inferred from genes detected in shotgun metagenomic sequencing). Our response variables consisted of these adult zebrafish gut microbiome sequencing samples, as well as four adult zebrafish behavior measures (free swim distance, shoaling nnd, shoaling iid, and shoaling speed). We used a Kolmogorov-Smirnov test to evaluate whether the distributions of behavioral metrics differed between control and experimental treatments. We used linear regressions to model alpha-diversity and beta-dispersion as the dependent variables. For beta-diversity, we optimized a dbRDA constrained ordination model then used the optimized model formula from this in a PERMANOVA to determine significance. Further, we included sex as a covariate in these regressions. We also identified biologically relevant ASVs and putative pathways that were indicative of BaP exposure, AHR2 morpholino treatment, or both by fitting relative abundance distributions to a compound Poisson generalized linear model with a log link function (MicroViz package [version 0.10.6] with phyloseq [version 1.42.0] objects as input) (Barnett, Arts, and Penders 2021; McMurdie and Holmes 2013).

We used R version 4.2.2 (2022-10-31) (R Core Team 2019) and associated packages to carry out statistical analyses and graphing. All code and associated files can be found on the project github page (https://github.com/aalexiev/ZebrafishBaPIntergenerationalMB). Graphs were produced using ggplot (version 3.4.1) (Wickham 2016) and the companion package ggpubr (version 0.6.0) (Alboukadel Kassambara 2023). Community diversity metrics and associated statistics were carried out with the vegan package (2.6-4) (Jari Oksanen et al. 2019).

Four behavior measures were analyzed first with the same approach from past zebrafish toxicology research (Knecht et al. 2017b; Reif et al. 2016), and secondarily using a previously used approach where area under the curve (AUC) was used in a linear model with treatment variables (Stagaman et al. 2024). Since the camera model was updated such that F0 has less-sensitive measurements than F1 and F2, we modeled and graphed each generation separately and compared metrics between treatments within each generation. With regard to the first approach, we produced a cumulative density distribution for each metric per treatment per generation (e.g., F0 AHR2Mo-/BaP-was one distribution, thus twelve total). Within each generation, we ran a Kolmogorov-Smirnov test comparing each treatment distribution (AHR2Mo-/BaP+, AHR2Mo+/BaP-, AHR2Mo+/BaP+) to the control (AHR2Mo-/BaP-), where a significant p-value indicates that the distributions are different from each other (Fig S1, Table S4). Regarding the second approach, we calculated the area under the curve (AUC) for each of four behavior metrics versus timepoint, per fish, within each generation (i.e., we filter the whole dataset to represent only F0 fish, then calculate AUC per fish on a graph of free swim distance versus timepoint, then pair this data with that fish’s metadata on what treatment each came from). We then ran a linear regression model for each generation with AUC as the independent variable and BaP exposure, AHR2 morpholino, and their interaction.

We first determined if there was an effect of BaP exposure on the gut microbiome of zebrafish and whether it was dependent on the AHR2 pathway using just F0 generation 16S and shotgun metagenomic sequence data. We tested several different gut microbiome alpha- and beta-diversity metrics, which are included in the supplement. We calculated these metrics for two datasets: ASVs and metabolic pathways encoded in the metagenome, derived from 16S and shotgun metagenomic sequence data, respectively. We then applied this question across generations by including sequence data from all generations in our models as well as analyzing F1 and F2 generations individually. For alpha-diversity, we report richness and Shannon diversity metrics (Shannon 1948) in the main text, as these are exemplar metrics that represent the overarching trends observed across our analyses, but also evaluated Simpson metrics (Simpson 1949) for all treatments and generations in the initial exploration of the data (Simpson metrics gave congruent results to Shannon). Similarly, for beta-diversity, we report Bray-Curtis dissimilarity (Bray and Curtis 1957) in the main text, but also evaluated Jaccard (Jaccard 1912)and Sørenson (Dice 1945; Sørensen 1948) metrics, which gave the same results. For alpha-diversity metrics and beta-dispersion, we ran linear regression models using AIC to choose the optimal model. For beta-diversity metrics, dbRDA was used to graph the ordination and ordistep to choose the optimal model to constrain upon. We then used this optimal model from the ordination as the input to a PERMANOVA model to determine the significance and area of effect of the covariates.

We wanted to visualize how beta-diversity varied across the three generations within each treatment, so we created a Bray-Curtis based metric to evaluate this. We first calculated Bray-Curtis dissimilarity for the full dataset. Then, we calculated the difference in dissimilarity between the F0 centroid all points in the F0 generation (as a baseline), F1, and F2. We then normalized to the “F0 centroid to F0 points” distance so that all distances for F1 and F2 were relative to F0, and F0 was on average zero. For example, the F0 metric was calculated as: distance from centroid F0 to each point in F0 divided by median distance from centroid F0 to each point in F0, minus 1. The calculation for F1 was distance from centroid F0 to each point in F1 divided by median distance from centroid F0 to each point in F0, minus 1. This represents, essentially, the dissimilarity in microbiomes between each of F1 and F2 compared to F0. We did this for each exposure/morpholino treatment individually. We use this measure to compare gut microbiome beta-diversity trends across generations.

Subsequently, indicator analyses were performed with the MicroViz package (version 0.10.6) (Barnett, Arts, and Penders 2021), using a compound Poisson generalized linear model to fit each taxon or pathway abundance distribution to BaP exposure, AHR2 morpholino status, or the combination of both. We used the incidence-rate ratio (IRR) to represent the degree and direction of interaction and graphed the log(IRR) for ease of interpretation (i.e., a positive value is an increased ASV/pathway relative abundance associated with that variable). Significance was determined via Wald test and p-values were FDR-adjusted (alpha = 0.05).

## Supplemental Figures and Tables

**Supplemental Figure 1:**
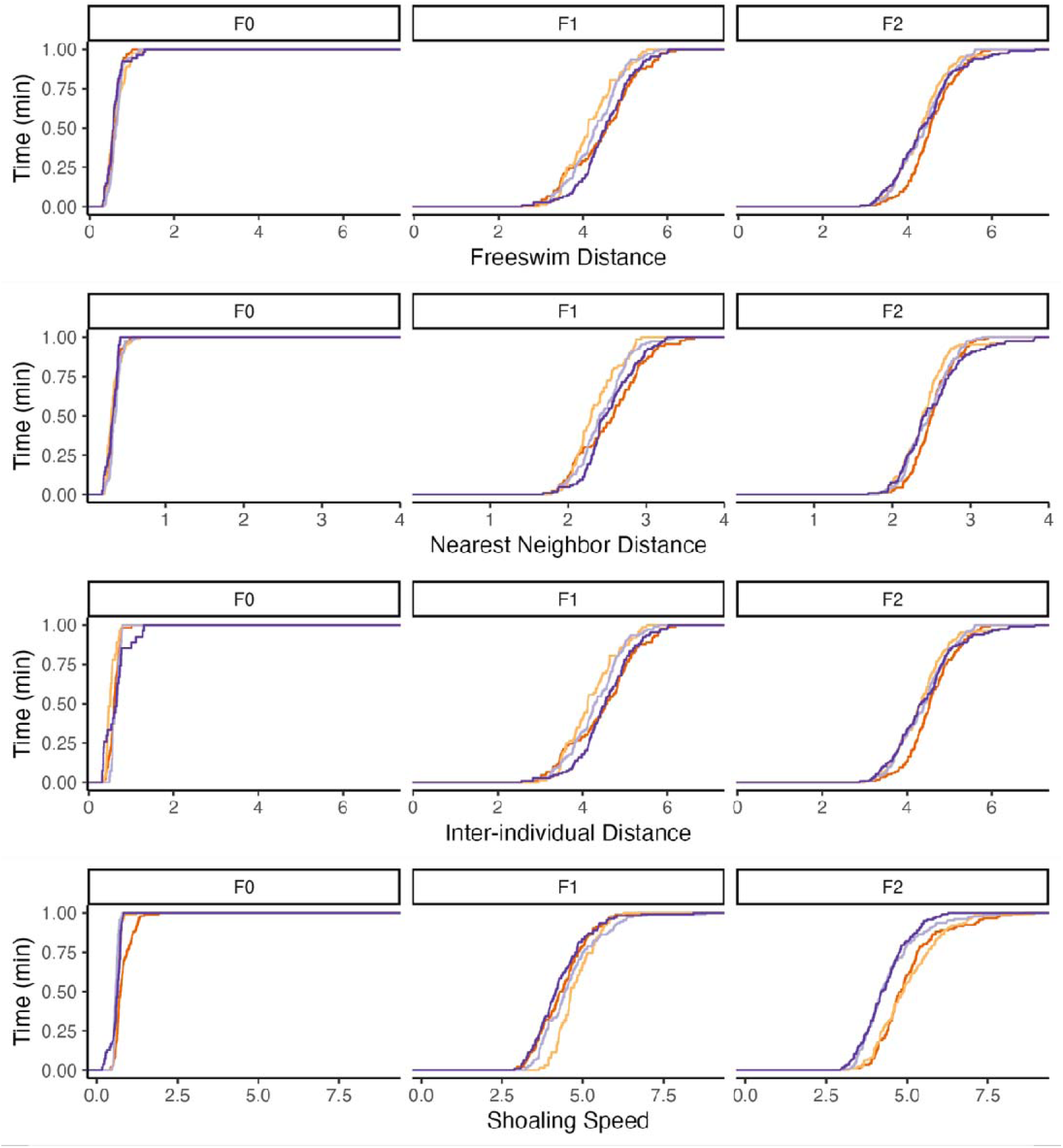
CDF distributions of four behavioral measures. Each of four behavior measures are shown per row, while the generations are shown per column. Each color represents one of four treatments; the KS test was run between the control treatment (AHR2Mo-/ BaP-, the dark purple line) and the other treatments. See Figure 7 and Table S4 for which statistical comparisons were significant.

**Supplemental Figure 2:**
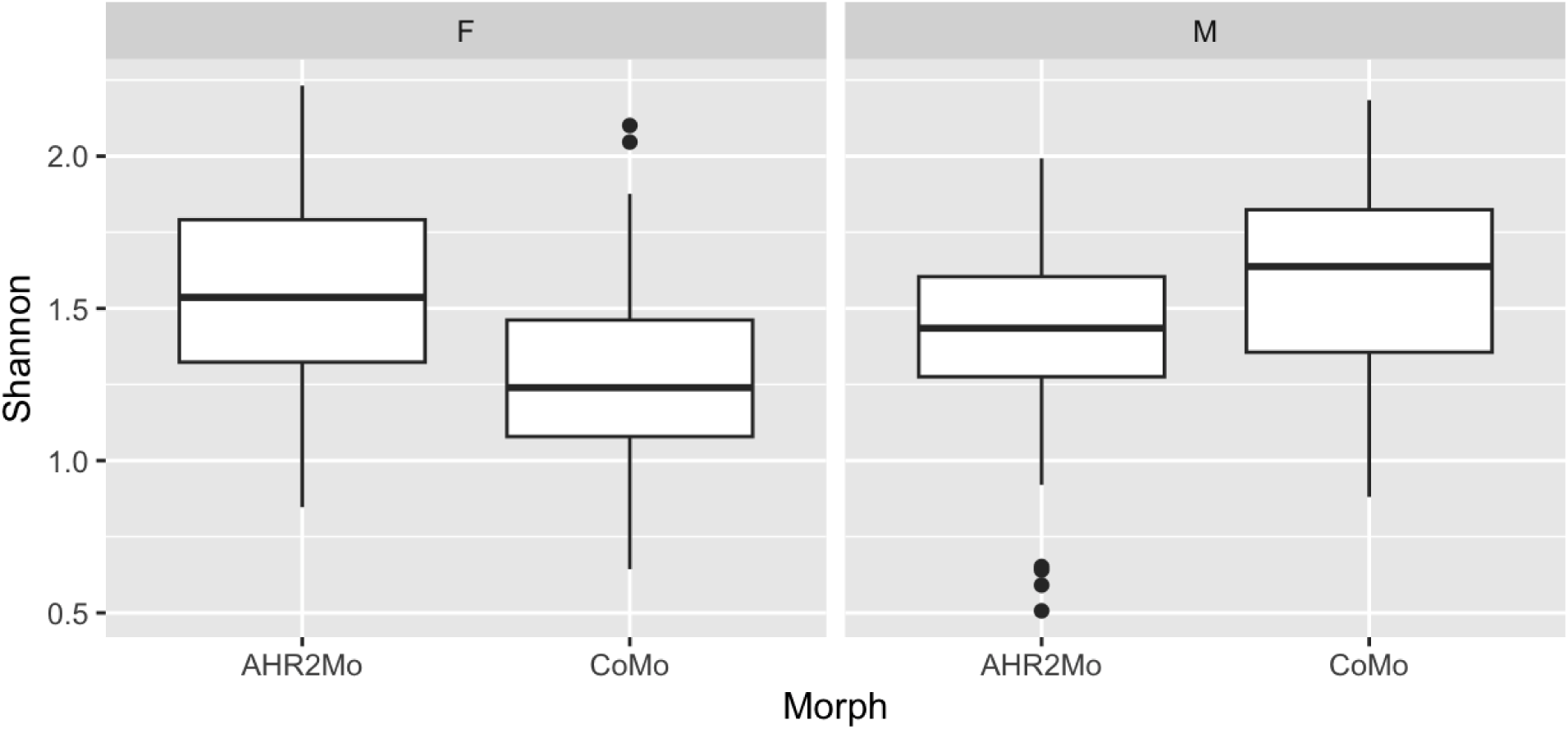
Box and whisker plot of taxonomic Shannon diversity. The plot is facet wrapped by sex of the fish and the morpholino type is shown along the x-axis. Sex (F for female, M for male) and morpholino status (AHR2Mo representing AHR2 morpholino treatment and CoMo representing the control morpholino) were the two covariates retained in the optimal linear regression model.

**Supplemental Figure 3:**
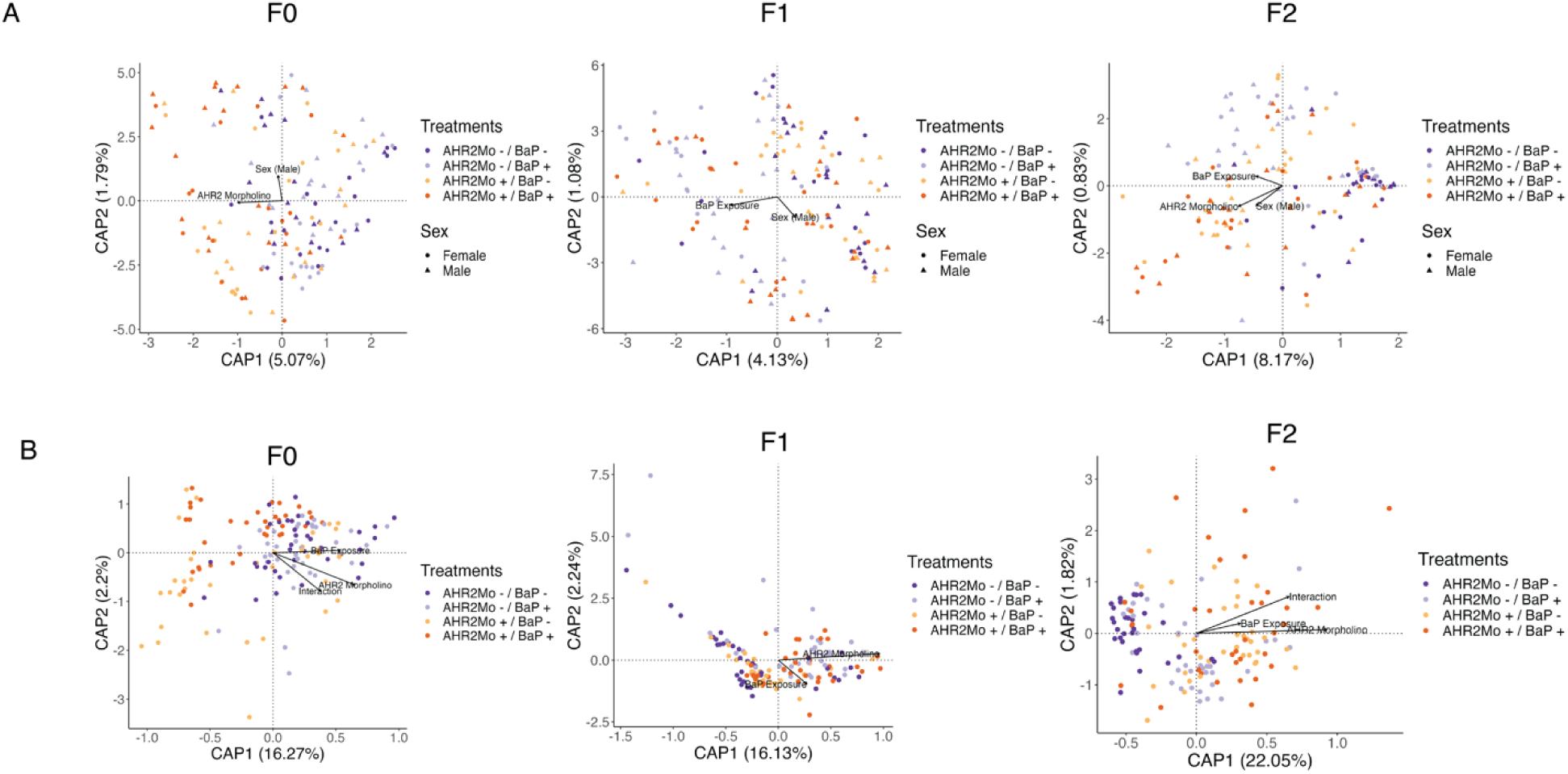
(A) dbRDA ordinations showing the differences between each generation of zebrafish using the 16S data. Each plot is labeled at the top with the corresponding generation. The colors show treatments and shapes show sex. (B) likewise, the shotgun metagenomic data dbRDA ordinations per generation with the colors showing treatments.

**Table S1:** Statistical outcomes from behavioral analysis, using the KS test to compare the CDF of the control group (AHR2Mo-/BaP-) to each treatment group.

| Behavior metric | Generation | Experimental group (compared to control) | KS test statistic | P-value (alpha $\leq 0.05$) | significance (asterisk is significant) |
| --- | --- | --- | --- | --- | --- |
| Freeswim distance (cm) | F0 | AhR2Mo - / BaP + | 0.2407407 | 0.085776 | ns |
|  | F0 | AhR2Mo + / BaP - | 0.1728395 | 0.2593598 | ns |
|  | F0 | AhR2Mo + / BaP + | 0.2296296 | 0.0460685 | * |
|  | F1 | AhR2Mo - / BaP + | 0.1759259 | 0.0706873 | ns |
|  | F1 | AhR2Mo + / BaP - | 0.3333333 | 0.0005538 | * |
|  | F1 | AhR2Mo + / BaP + | 0.1703704 | 0.0994281 | ns |
|  | F2 | AhR2Mo - / BaP + | 0.1111111 | 0.4694025 | ns |
|  | F2 | AhR2Mo + / BaP - | 0.1239316 | 0.3542932 | ns |
|  | F2 | AhR2Mo + / BaP + | 0.2336182 | 0.0043525 | * |
| Shoaling nearest-neighbor distance (nnd) (cm) | F0 | AhR2Mo - / BaP + | 0.2222222 | 0.0306982 | * |
|  | F0 | AhR2Mo + / BaP - | 0.2716049 | 0.0038011 | * |
|  | F0 | AhR2Mo + / BaP + | 0.1296296 | 0.3612486 | ns |
|  | F1 | AhR2Mo - / BaP + | 0.1481481 | 0.1867397 | ns |
|  | F1 | AhR2Mo + / BaP - | 0.3240741 | 0.0008351 | * |
|  | F1 | AhR2Mo + / BaP + | 0.1888889 | 0.0497375 | * |
|  | F2 | AhR2Mo - / BaP + | 0.0873016 | 0.7442496 | ns |
|  | F2 | AhR2Mo + / BaP - | 0.2307692 | 0.0039359 | * |
|  | F2 | AhR2Mo + / BaP + | 0.1794872 | 0.046139 | * |
| Shoaling interindividual distance (iid) (cm) | F0 | AhR2Mo - / BaP + | 0.3333333 | 0.0995625 | ns |
|  | F0 | AhR2Mo + / BaP - | 0.4074074 | 0.0206395 | * |
|  | F0 | AhR2Mo + / BaP + | 0.2592593 | 0.1684132 | ns |
|  | F1 | AhR2Mo - / BaP + | 0.1759259 | 0.0706873 | ns |
|  | F1 | AhR2Mo + / BaP - | 0.3333333 | 0.0005538 | * |
|  | F1 | AhR2Mo + / BaP + | 0.1703704 | 0.0994281 | ns |
|  | F2 | AhR2Mo - / BaP + | 0.1111111 | 0.4694025 | ns |
|  | F2 | AhR2Mo + / BaP - | 0.1239316 | 0.3542932 | ns |
|  | F2 | AhR2Mo + / BaP + | 0.2336182 | 0.0043525 | * |
| Shoaling speed (cm/min) | F0 | AhR2Mo - / BaP + | 0.2925926 | 0.0230448 | * |
|  | F0 | AhR2Mo + / BaP - | 0.1728395 | 0.2511765 | ns |
|  | F0 | AhR2Mo + / BaP + | 0.3838384 | 0.0000364 | * |
|  | F1 | AhR2Mo - / BaP + | 0.212963 | 0.0149208 | * |
|  | F1 | AhR2Mo + / BaP - | 0.398148<br>1 | 0.0000037 | * |
|  | F1 | AhR2Mo + / BaP + | 0.109259<br>3 | 0.556501 | ns |
|  | F2 | AhR2Mo - / BaP + | 0.080586<br>1 | 0.825658 | ns |
|  | F2 | AhR2Mo + / BaP - | 0.315628<br>8 | 0.0000113 | * |
|  | F2 | AhR2Mo + / BaP + | 0.324786<br>3 | 0.0000087 | * |

**Table S2:** All taxonomic (16S) models used to test whether microbial taxonomic diversity metrics were associated with different experimental treatment variables.

| Metric | Input dataset | Optimal mathematical model with formula | adjusted r-squared | output |
| --- | --- | --- | --- | --- |
| Alpha diversity | F0 | Linear regression model:<br>Taxonomic Richness ~ Exposure + Morpholino + Sex + Exposure:Morpholino + Sex:Exposure + Sex:Morpholino | 0.1661 | Estimate Std. Error t value Pr(>\|t\|)<br>(Intercept) 20.0161 1.3637 14.677 < 2e-16 ***<br>ExposureBaP -0.9774 1.8147 -0.539 0.59102<br>MorpholinoAHR2Mo 3.0205 1.7975 1.680 0.09520 .<br>SexM 1.7307 1.8331 0.944 0.34679<br>ExposureBaP:MorpholinoAHR2Mo -6.7901 2.1286 -3.190 0.00177 **<br>ExposureBaP:SexM 3.8957 2.1288 1.830 0.06946 .<br>MorpholinoAHR2Mo:SexM -5.3107 2.1288 -2.495 0.01381 * |
|  | F0 | Linear regression model:<br>Taxonomic Shannon diversity ~ Exposure + Morpholino + Sex + Exposure:Morpholino + Morpholino:Sex | 0.09818 | Estimate Std. Error t value Pr(>\|t\|)<br>(Intercept) 1.31062 0.06928 18.917 < 2e-16 ***<br>ExposureBaP -0.05531 0.08187 -0.676 0.500413<br>MorpholinoAHR2Mo 0.17244 0.09804 1.759 0.080842 .<br>SexM 0.30530 0.08193 3.726 0.000284 ***<br>ExposureBaP:MorpholinoAHR2Mo 0.17864 0.11605 1.539 0.126057<br>MorpholinoAHR2Mo:SexM -0.47104 0.11610 -4.057 8.33e-05 *** |
|  | F1 | Linear regression model:<br>Taxonomic Richness ~ Morpholino + Sex | 0.06068 | Estimate Std. Error t value Pr(>\|t\|)<br>(Intercept) 20.853 2.639 7.901 7.61e-13 ***<br>MorpholinoAHR2Mo -8.570 3.034 -2.824 0.00543 **<br>SexM 5.125 3.034 1.689 0.09344 . |
|  | F1 | Linear regression model:<br>Taxonomic Shannon diversity ~ Morpholino + Sex | 0.07838 | Estimate Std. Error t value Pr(>\|t\|)<br>(Intercept) 1.49776 0.05848 25.610 < 2e-16 ***<br>MorpholinoAHR2Mo -0.19131 0.06724 -2.845 0.00511 **<br>SexM 0.15778 0.06723 2.347 0.02034 * |
|  | F2 | Linear regression model:<br>Taxonomic Richness ~ Exposure + Morpholino | 0.2197 | Estimate Std. Error t value Pr(>\|t\|)<br>(Intercept) 14.1933 0.9331 15.211 < 2e-16 ***<br>ExposureBaP 3.2774 1.0432 3.142 0.00207 **<br>MorpholinoAHR2Mo 5.8346 1.0444 5.587 1.24e-07 *** |
|  | F2 | Linear regression model:<br>Taxonomic Shannon diversity ~ Exposure + Morpholino + Sex + Exposure:Morpholino | 0.06875 | Estimate Std. Error t value Pr(>\|t\|)<br>(Intercept) 1.32257 0.06409 20.638 < 2e-16 ***<br>ExposureBaP 0.25454 0.08144 3.125 0.00218 **<br>MorpholinoAHR2Mo 0.05061 0.08036 0.630 0.52988<br>SexM 0.08760 0.05607 1.562 0.12057<br>ExposureBaP:MorpholinoAHR2Mo -0.22227 0.11226 -1.980 0.04979 * |
|  | Whole dataset | Linear regression model:<br>Taxonomic Richness ~<br>Generation + Exposure +<br>Morpholino + Sex +<br>Generation:Exposure +<br>Generation:Morpholino +<br>Exposure:Morpholino +<br>Morpholino:Sex | 0.07561 | Estimate Std. Error t value Pr(>\|t\|)<br>(Intercept) 19.3848 1.9767 9.807 < 2e-16 ***<br>GenerationF1 0.5712 2.4408 0.234 0.815073<br>GenerationF2 -8.4352 2.5117 -3.358 0.000858 ***<br>ExposureBaP -0.7502 2.3159 -0.324 0.746152<br>MorpholinoAHR2Mo 1.1709 2.5653 0.456 0.648305<br>SexM 4.5570 1.6558 2.752 0.006186 **<br>GenerationF1:ExposureBaP 3.5157 2.8317 1.242 0.215108<br>GenerationF2:ExposureBaP 5.8387 2.8592 2.042 0.041783 *<br>GenerationF1:MorpholinoAHR2Mo -6.0585 2.8319 -2.139 0.032997 *<br>GenerationF2:MorpholinoAHR2Mo 8.8692 2.8589 3.102 0.002053 **<br>ExposureBaP:MorpholinoAHR2Mo -3.3254 2.3282 -1.428 0.153969<br>MorpholinoAHR2Mo:SexM -4.6052 2.3286 -1.978 0.048631 * |
|  | Whole dataset | Linear regression model:<br>Taxonomic Shannon diversity ~<br>Generation + Exposure +<br>Morpholino + Sex +<br>Generation:Morpholino +<br>Morpholino:Sex | 0.04942 | Estimate Std. Error t value Pr(>\|t\|)<br>(Intercept) 1.30500 0.05239 24.912 < 2e-16 ***<br>GenerationF1 0.14370 0.06095 2.358 0.018846 *<br>GenerationF2 0.06806 0.06254 1.088 0.277104<br>ExposureBaP 0.06882 0.03554 1.937 0.053474 .<br>MorpholinoAHR2Mo 0.11124 0.07046 1.579 0.115142<br>SexM 0.18503 0.05056 3.660 0.000285 ***<br>GenerationF1:MorpholinoAHR2Mo -0.22229 0.08648 -2.570 0.010511 *<br>GenerationF2:MorpholinoAHR2Mo -0.09791 0.08731 -1.121 0.262743<br>MorpholinoAHR2Mo:SexM -0.15876 0.07105 -2.235 0.025980 * |
| Beta diversity | F0 | PERMANOVA using optimized model formula from dbRDA:<br>Taxonomic Bray-Curtis Distance Matrix ~ Morpholino + Sex | NA | Df SumOfSqs R2 F Pr(>F)<br>Morpholino 1 1.3780 0.06135 9.2937 0.001 ***<br>Sex 1 0.4732 0.02107 3.1912 0.028 *<br>Residual 139 20.6102 0.91758<br>Total 141 22.4614 1.00000 |
|  | F1 | PERMANOVA using optimized model formula from dbRDA:<br>Taxonomic Bray-Curtis Distance Matrix ~ Exposure + Sex | NA | Df SumOfSqs R2 F Pr(>F)<br>Exposure 1 1.0658 0.04367 6.4650 0.001 ***<br>Sex 1 0.4256 0.01744 2.5815 0.062 .<br>Residual 139 22.9149 0.93889<br>Total 141 24.4063 1.00000 |
|  | F2 | PERMANOVA using optimized model formula from dbRDA:<br>Taxonomic Bray-Curtis Distance Matrix ~ Morpholino + Exposure + Morpholino:Exposure | NA | Df SumOfSqs R2 F Pr(>F)<br>Morpholino 1 1.3338 0.06051 9.0837 0.001 ***<br>Exposure 1 0.5963 0.02705 4.0611 0.005 **<br>Morpholino:Exposure 1 0.5847 0.02652 3.9821 0.015 *<br>Residual 133 19.5289 0.88592<br>Total 136 22.0437 1.00000 |
|  | Whole dataset | PERMANOVA using optimized model formula from dbRDA:<br>Taxonomic Bray-Curtis Distance Matrix ~ Generation + Exposure + Sex + Generation:Exposure | NA | Df SumOfSqs R2 F Pr(>F)<br>Generation 2 2.304 0.03233 7.1571 0.001 ***<br>Exposure 1 0.835 0.01171 5.1851 0.004 **<br>Sex 1 0.484 0.00680 3.0084 0.023 *<br>Generation:Exposure 2 1.001 0.01405 3.1092 0.007 **<br>Residual 414 66.635 0.93512<br>Total 420 71.258 1.00000 |
| Beta dispersion | F0 | Linear regression model:<br>Taxonomic Beta-dispersion<br>distances ~ Exposure +<br>Morpholino + Sex +<br>Exposure:Morpholino +<br>Sex:Exposure + Sex:Morpholino | 0.04192 | Estimate Std. Error t value Pr(>\|t\|)<br>(Intercept) 0.33874 0.02249 15.062 <2e-16 ***<br>MorpholinoAHR2Mo 0.01301 0.03226 0.403 0.6874<br>SexM -0.01798 0.03250 -0.553 0.5809<br>MorpholinoAHR2Mo:SexM 0.07943 0.04594 1.729<br>0.0861 . |
|  | F1 | Linear regression model:<br>Taxonomic Beta-dispersion<br>distances ~ Exposure +<br>Morpholino +<br>Exposure:Morpholino | -0.003852 | Estimate Std. Error t value Pr(>\|t\|)<br>(Intercept) 0.39873 0.02450 16.277 <2e-16 ***<br>ExposureBaP -0.04656 0.03464 -1.344 0.181<br>MorpholinoAHR2Mo -0.03777 0.03515 -1.075<br>0.284<br>ExposureBaP:MorpholinoAHR2Mo 0.07495 0.04935<br>1.519 0.131 |
|  | F2 | Linear regression model:<br>Taxonomic Beta-dispersion<br>distances ~ Morpholino + Sex +<br>Morpholino:Sex | 0.07738 | Estimate Std. Error t value Pr(>\|t\|)<br>(Intercept) 0.33132 0.02408 13.761 <2e-16 ***<br>MorpholinoAHR2Mo 0.09495 0.03357 2.828 0.0054<br>**<br>SexM -0.02255 0.03486 -0.647 0.5188<br>MorpholinoAHR2Mo:SexM -0.07063 0.04807 -1.469<br>0.1441 |
|  | Whole dataset | Linear regression model:<br>Taxonomic Beta-dispersion<br>distances ~ Morpholino | 0.01346 | Estimate Std. Error t value Pr(>\|t\|)<br>(Intercept) 0.343350 0.009875 34.770 < 2e-16 ***<br>MorpholinoAHR2Mo 0.036013 0.013883 2.594 0.00982 ** |

**Table S3:** All pathway (metagenomic) models used to test whether microbially derived pathway diversity metrics were associated with different experimental treatment variables.

| Input dataset | Optimal mathematical model with formula | adjusted r-squared | output |
| --- | --- | --- | --- |
| F0 | Linear regression model: Pathway Richness ~ Exposure + Morpholino + Sex + Exposure:Morpholino + Morpholino:Sex | 0.02051 | Estimate Std. Error t value Pr(>\|t\|)<br>(Intercept) 32.6784 1.3933 23.453 <2e-16 ***<br>ExposureBaP -2.7778 1.6379 -1.696 0.0921 .<br>MorpholinoAHR2Mo 0.5985 1.9712 0.304 0.7619<br>SexM 3.6223 1.6404 2.208 0.0289 *<br>ExposureBaP:MorpholinoAHR2Mo 3.2972 2.3204 1.421 0.1576<br>MorpholinoAHR2Mo:SexM -3.8556 2.3226 -1.660 0.0992 . |
| F0 | Linear regression model: Pathway Shannon diversity ~ Morpholino + Sex + Morpholino:Sex | 0.04728 | Estimate Std. Error t value Pr(>\|t\|)<br>(Intercept) 4.52135 0.01192 379.315 <2e-16 ***<br>MorpholinoAHR2Mo -0.01160 0.01721 -0.674 0.5014<br>SexM 0.02969 0.01735 1.711 0.0892 .<br>MorpholinoAHR2Mo:SexM -0.04152 0.02452 -1.693 0.0926 . |
| F1 | Linear regression model: Pathway Richness ~ Exposure + Morpholino + Sex + Exposure:Morpholino | 0.02326 | Estimate Std. Error t value Pr(>\|t\|)<br>(Intercept) 33.790 1.365 24.757 <2e-16 ***<br>ExposureBaP -1.371 1.727 -0.794 0.4286<br>MorpholinoAHR2Mo -1.643 1.763 -0.932 0.3531<br>SexM 2.259 1.225 1.844 0.0673 .<br>ExposureBaP:MorpholinoAHR2Mo 4.400 2.450 1.796 0.0747 . |
| F1 | Linear regression model: Pathway Shannon diversity ~ Exposure + Morpholino + Exposure:Morpholino | 0.09264 | Estimate Std. Error t value Pr(>\|t\|)<br>(Intercept) 4.4736106 0.0130752 342.145 < 2e-16 ***<br>ExposureBaP 0.0650879 0.0182325 3.570 0.000495 ***<br>MorpholinoAHR2Mo 0.0001005 0.0186307 0.005 0.995705<br>ExposureBaP:MorpholinoAHR2Mo -0.0391796 0.0258849 -1.514 0.132463 |
| F2 | Linear regression model: Pathway Richness ~ Morpholino + Sex + Morpholino:Sex | 0.02107 | Estimate Std. Error t value Pr(>\|t\|)<br>(Intercept) 33.514 1.160 28.898 <2e-16 ***<br>MorpholinoAHR2Mo 1.736 1.629 1.066 0.2885<br>SexM 2.673 1.678 1.593 0.1135<br>MorpholinoAHR2Mo:SexM -5.395 2.331 -2.315 0.0221 * |
| F2 | Linear regression model: Pathway Shannon diversity ~ Exposure + Morpholino + Exposure:Morpholino | 0.1231 | Estimate Std. Error t value Pr(>\|t\|)<br>(Intercept) 4.45502 0.01147 388.345 < 2e-16 ***<br>ExposureBaP 0.06022 0.01565 3.848 0.000183 ***<br>MorpholinoAHR2Mo 0.05949 0.01565 3.801 0.000217 ***<br>ExposureBaP:MorpholinoAHR2Mo -0.05528 0.02172 -2.546 0.012027 * |
| Whole dataset | Linear regression model: Pathway Richness ~ Exposure + Morpholino + Sex + Exposure:Morpholino | 0.005263 | Estimate Std. Error t value Pr(>\|t\|)<br>(Intercept) 33.7658 0.7736 43.648 <2e-16 ***<br>ExposureBaP -0.7708 0.9795 -0.787 0.4318<br>MorpholinoAHR2Mo -1.0870 0.9861 -1.102 0.2709<br>SexM 1.2692 0.6894 1.841 0.0663 .<br>ExposureBaP:MorpholinoAHR2Mo 2.1932 1.3780 1.592 0.1123 |
| Whole dataset | Linear regression model: Pathway Shannon diversity ~ Generation + Exposure + Morpholino + Sex + Generation:Exposure + Generation:Morpholino + Exposure:Morpholino | 0.095 | Estimate Std. Error t value Pr(>\|t\|)<br>(Intercept) 4.517379 0.011329 398.755 < 2e-16 ***<br>GenerationF1 -0.048434 0.014773 -3.279 0.00113 **<br>GenerationF2 -0.062332 0.015008 -4.153 3.99e-05 ***<br>ExposureBaP 0.026185 0.013797 1.898 0.05842 .<br>MorpholinoAHR2Mo -0.013178 0.013797 -0.955 0.34010<br>SexM 0.010367 0.006964 1.489 0.13734<br>GenerationF1:ExposureBaP 0.037897 0.016997 2.230 0.02632 *<br>GenerationF2:ExposureBaP 0.024775 0.017006 1.457 0.14592<br>GenerationF1:MorpholinoAHR2Mo 0.012699 0.016997 0.747 0.45543<br>GenerationF2:MorpholinoAHR2Mo 0.063407 0.017006 3.729 0.00022 ***<br>ExposureBaP:MorpholinoAHR2Mo -0.037911 0.013925 -2.722 0.00676 ** |
| F0 | PERMANOVA using optimized model formula from dbRDA: Pathway Bray-Curtis Distance Matrix ~ Morpholino + Exposure + Morpholino:Exposure | NA | Df SumOfSqs R2 F Pr(>F)<br>Morpholino 1 0.14303 0.09031 16.163 0.001 ***<br>Exposure 1 0.09500 0.05999 10.736 0.001 ***<br>Morpholino:Exposure 1 0.10680 0.06744 12.070 0.001 ***<br>Residual 140 1.23883 0.78226<br>Total 143 1.58365 1.00000 |
| F1 | PERMANOVA using optimized model formula from dbRDA: Pathway Bray-Curtis Distance Matrix ~ Morpholino + Exposure | NA | Df SumOfSqs R2 F Pr(>F)<br>Morpholino 1 0.37431 0.16621 28.2680 0.001 ***<br>Exposure 1 0.07694 0.03416 5.8104 0.002 **<br>Residual 136 1.80085 0.79963<br>Total 138 2.25210 1.00000 |
| F2 | PERMANOVA using optimized model formula from dbRDA: Pathway Bray-Curtis Distance Matrix ~ Morpholino + Exposure + Morpholino:Exposure | NA | Df SumOfSqs R2 F Pr(>F)<br>Morpholino 1 0.32008 0.2131 39.3762 0.001 ***<br>Exposure 1 0.05017 0.0334 6.1716 0.002 **<br>Morpholino:Exposure 1 0.03439 0.0229 4.2313 0.004 **<br>Residual 135 1.09737 0.7306<br>Total 138 1.50201 1.0000 |
| Whole dataset | PERMANOVA using optimized model formula from dbRDA: Pathway Bray-Curtis Distance Matrix ~ Generation + Exposure + Morpholino + Generation:Exposure + Exposure:Morpholino + Generation:Morpholino | NA | Df SumOfSqs R2 F Pr(>F)<br>Generation 2 1.1709 0.21293 60.9419 0.001 ***<br>Exposure 1 0.1020 0.01856 10.6216 0.001 ***<br>Morpholino 1 0.0305 0.00555 3.1788 0.015 *<br>Generation:Exposure 2 0.0900 0.01636 4.6819 0.001 ***<br>Exposure:Morpholino 1 0.0664 0.01207 6.9086 0.001 ***<br>Generation:Morpholino 2 0.0811 0.01475 4.2222 0.001 ***<br>Residual 412 3.9579 0.71978<br>Total 421 5.4988 1.00000 |
| F0 | Linear regression model: Pathway Beta-dispersion distances ~ Morpholino | 0.01946 | Estimate Std. Error t value Pr(>\|t\|)<br>(Intercept) 0.076272 0.004971 15.344 <2e-16 ***<br>MorpholinoAHR2Mo 0.013772 0.007030 1.959 0.0521 . |
| F1 | Linear regression model: Pathway Beta-dispersion distances ~ Morpholino | 0.0185 | Estimate Std. Error t value Pr(>\|t\|)<br>(Intercept) 0.099326 0.008715 11.397 <2e-16 ***<br>MorpholinoAHR2Mo -0.023473 0.012369 -1.898 0.0598 . |
| F2 | Linear regression model: Pathway Beta-dispersion distances ~ Exposure + Morpholino | 0.06571 | Estimate Std. Error t value Pr(>\|t\|)<br>(Intercept) 0.060822 0.006345 9.585 <2e-16 ***<br>ExposureBaP 0.016833 0.007068 2.382 0.0186 *<br>MorpholinoAHR2Mo 0.017977 0.007068 2.543 0.0121 * |

| Whole dataset | Linear regression model: Pathway Beta-dispersion distances ~ Generation + Exposure + Morpholino + Generation:Exposure + Generation:Morpholino | 0.03911 | Estimate Std. Error t value Pr(>\|t\|) |
| --- | --- | --- | --- |
|  |  |  | (Intercept) 0.083441 0.008142 10.249 <2e-16 *** |
|  |  |  | GenerationF1 0.008263 0.011662 0.709 0.4790 |
|  |  |  | GenerationF2 -0.016293 0.011844 -1.376 0.1697 |
|  |  |  | ExposureBaP -0.017794 0.009401 -1.893 0.0591 . |
|  |  |  | MorpholinoAHR2Mo 0.002962 0.009401 0.315 0.7529 |
|  |  |  | GenerationF1:ExposureBaP 0.027597 0.013419 2.057 0.0404 * |
|  |  |  | GenerationF2:ExposureBaP 0.023332 0.013424 1.738 0.0829 . |
|  |  |  | GenerationF1:MorpholinoAHR2Mo -0.030512 0.013415 -2.275 0.0234 * |
|  |  |  | GenerationF2:MorpholinoAHR2Mo 0.011703 0.013424 0.872 0.3838 |

**Table S4:**
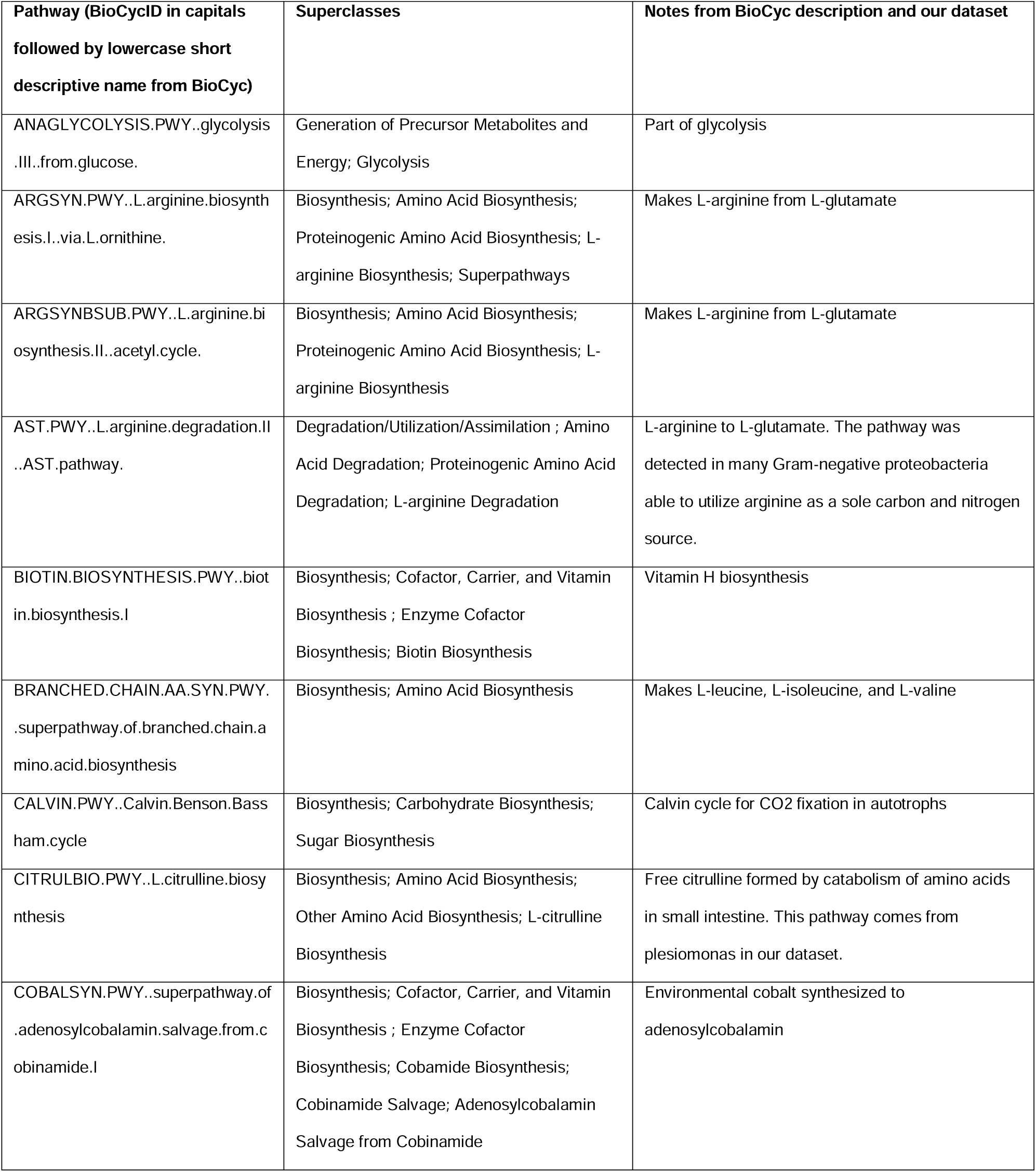
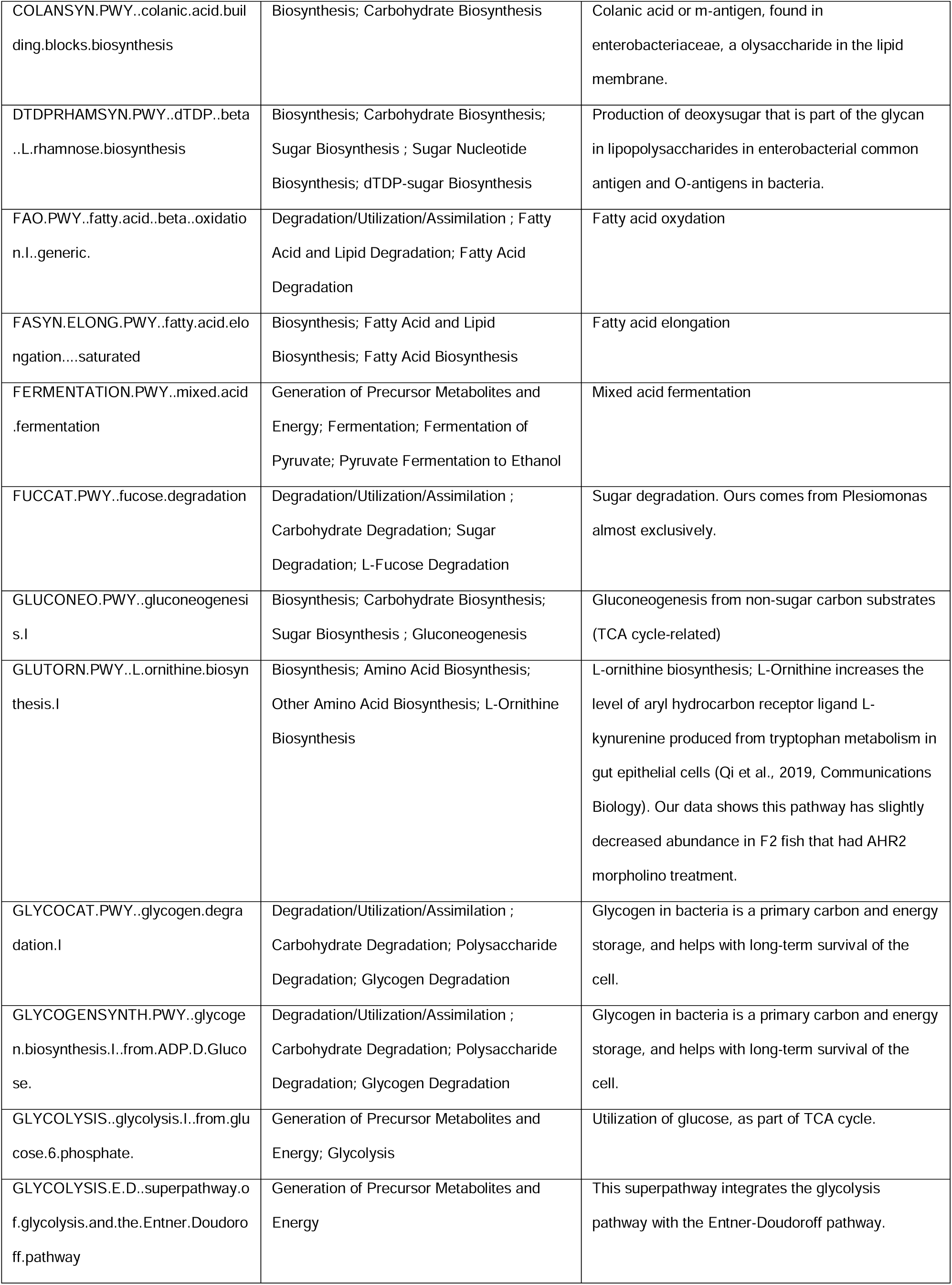
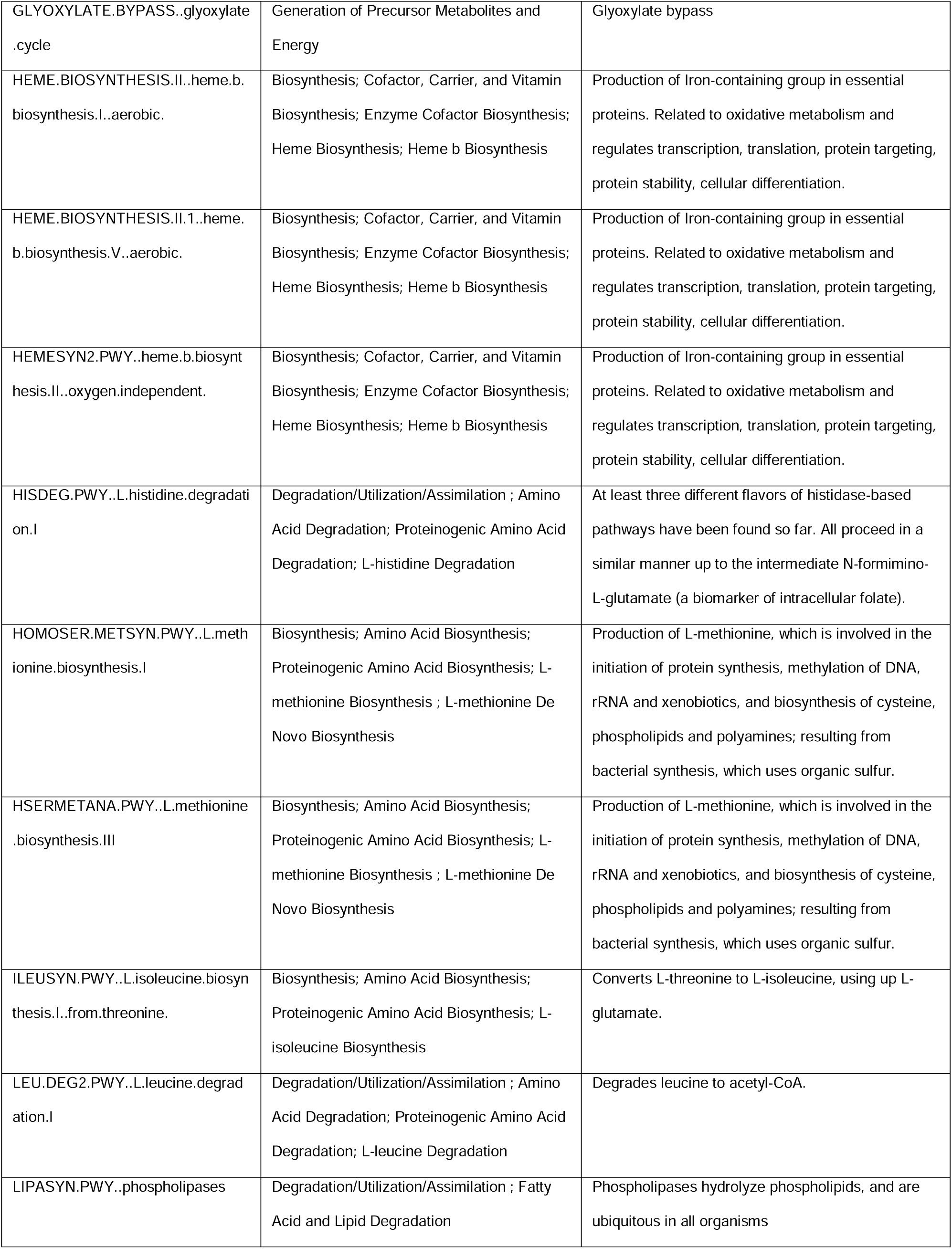
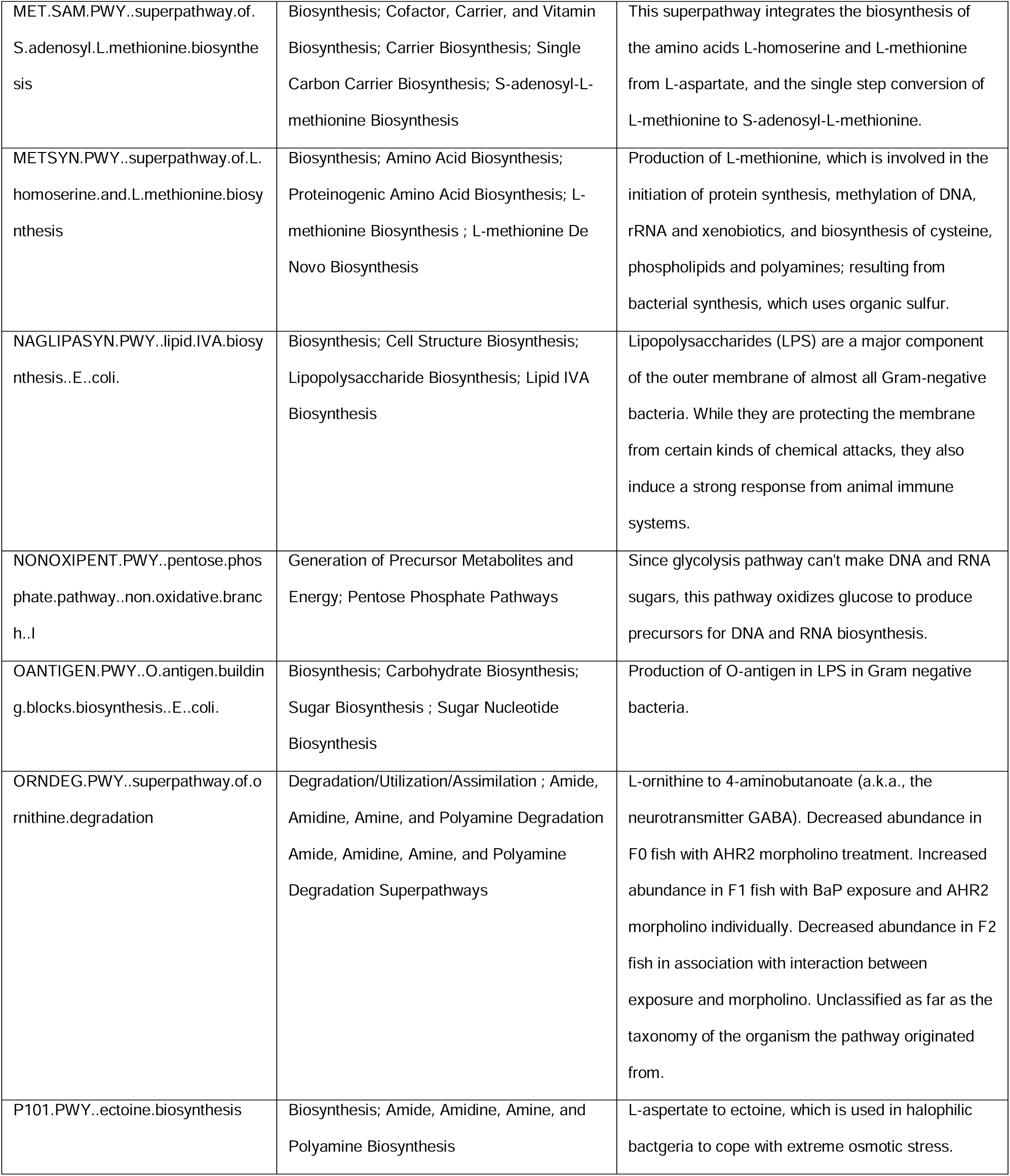
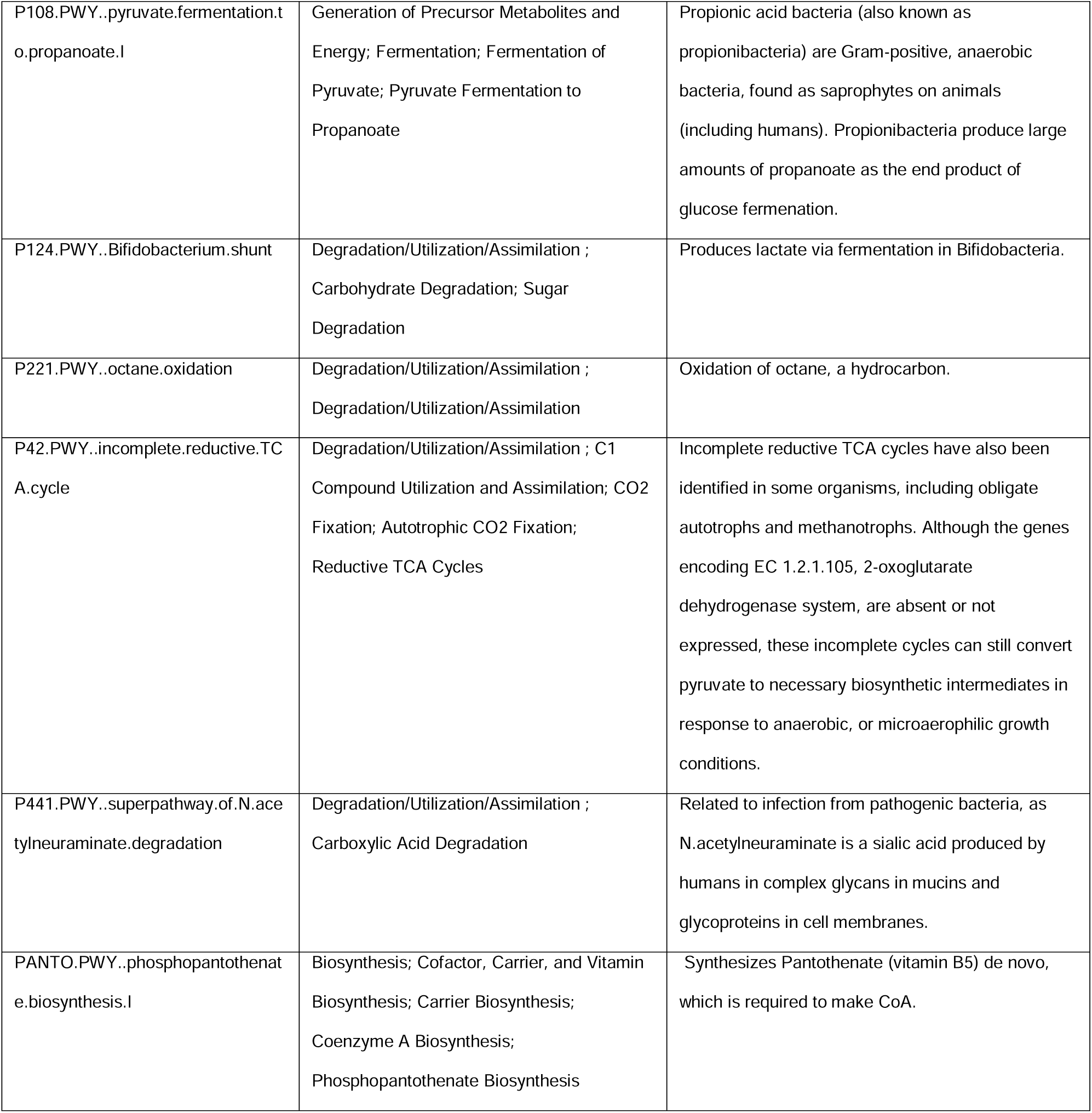
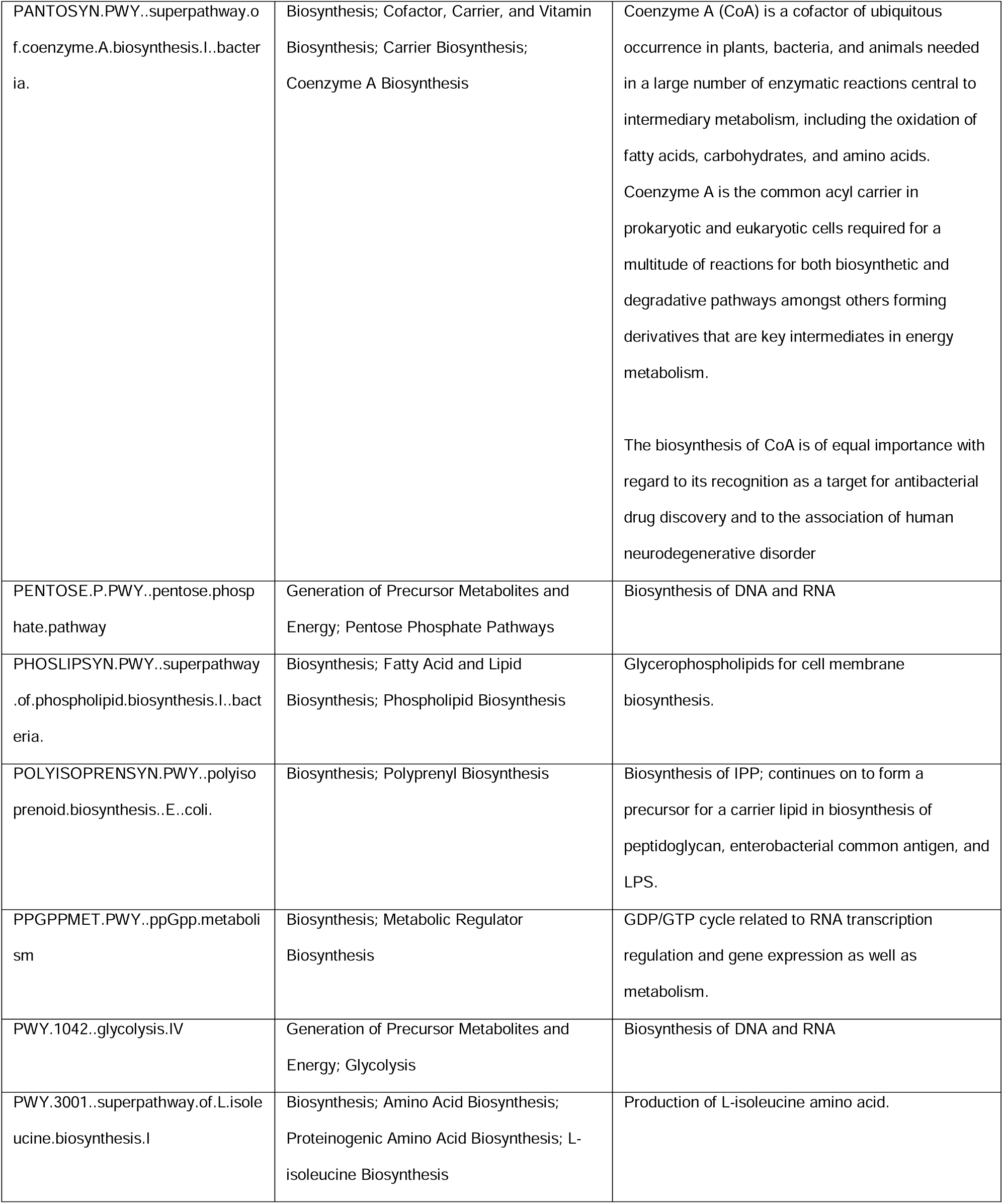
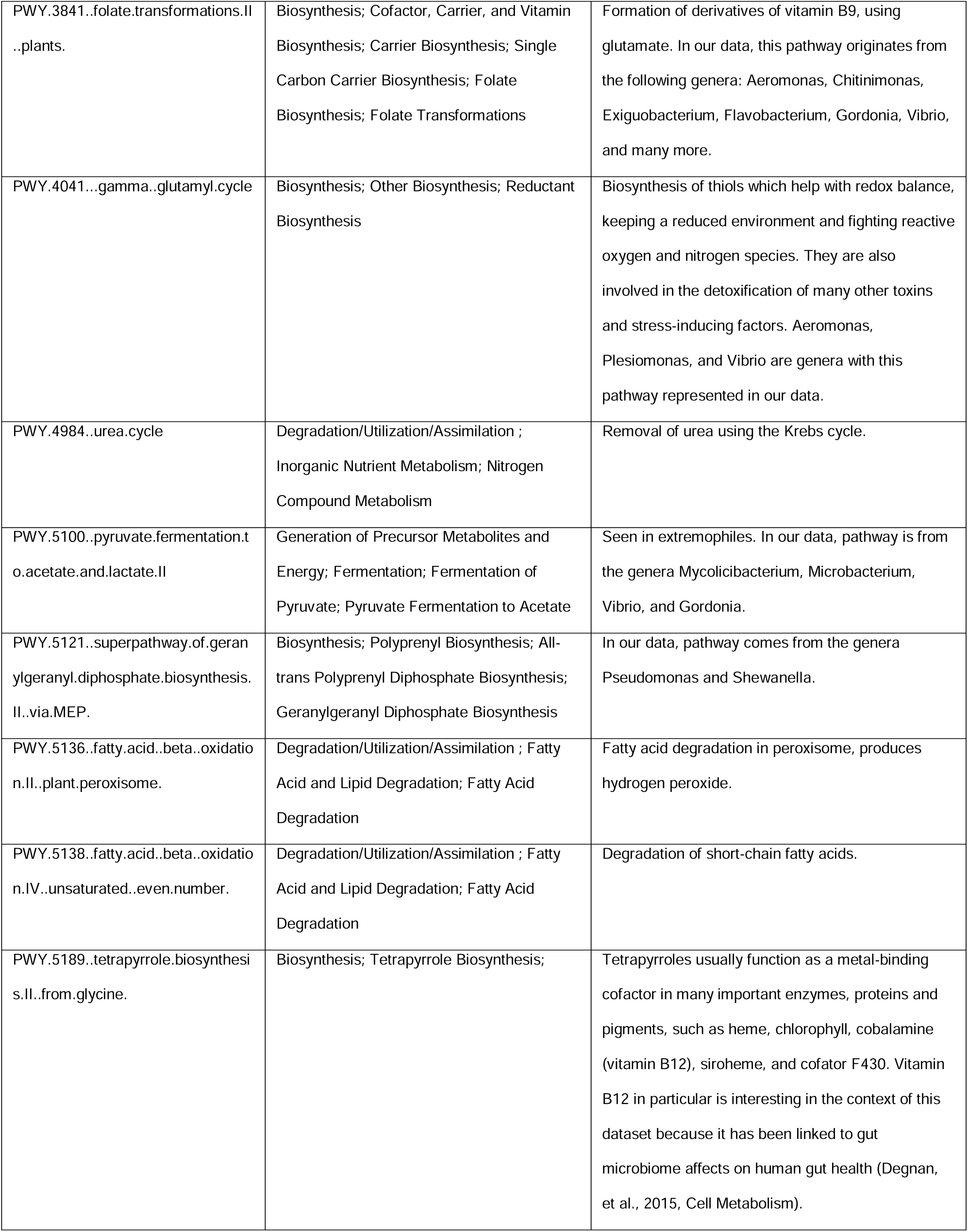
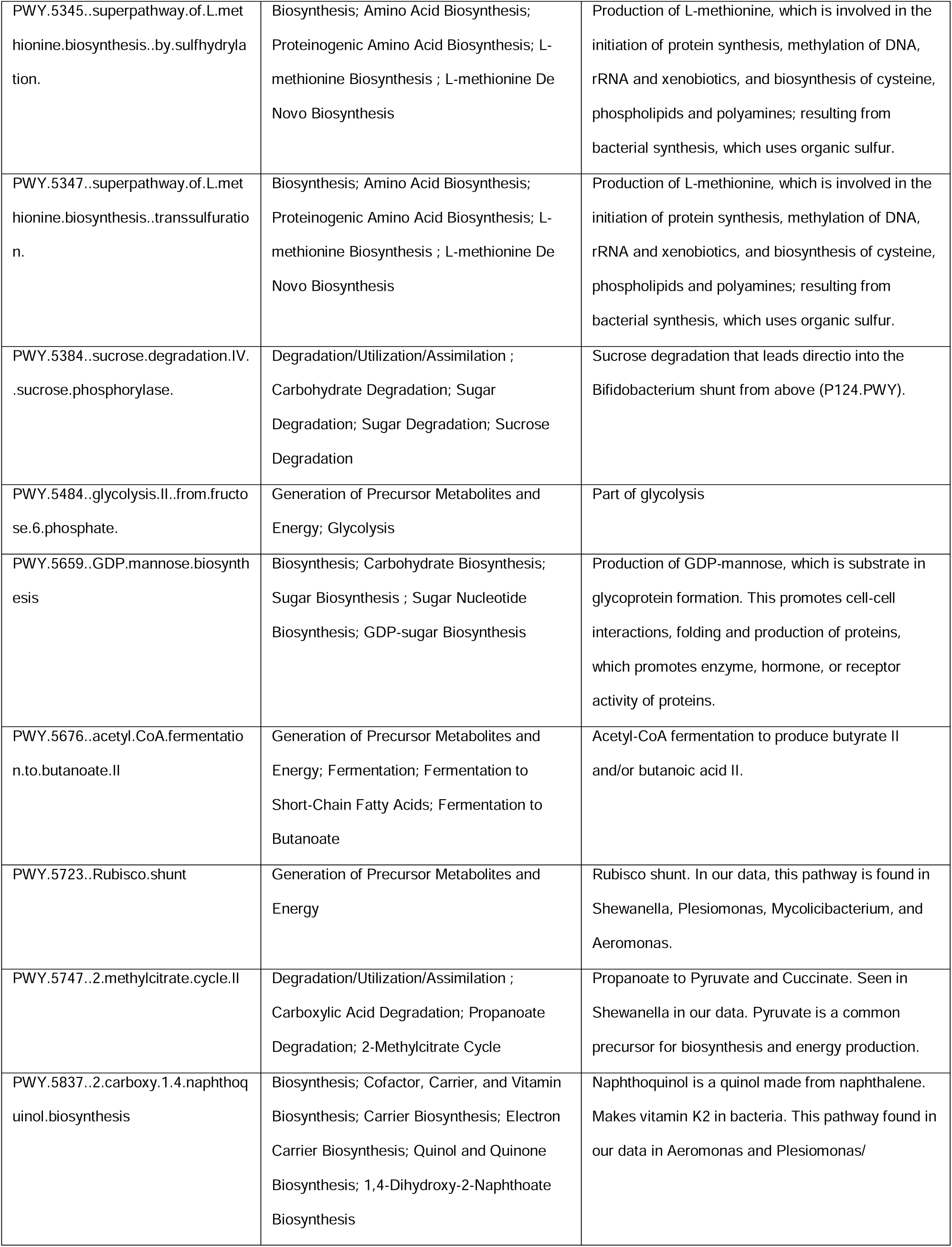
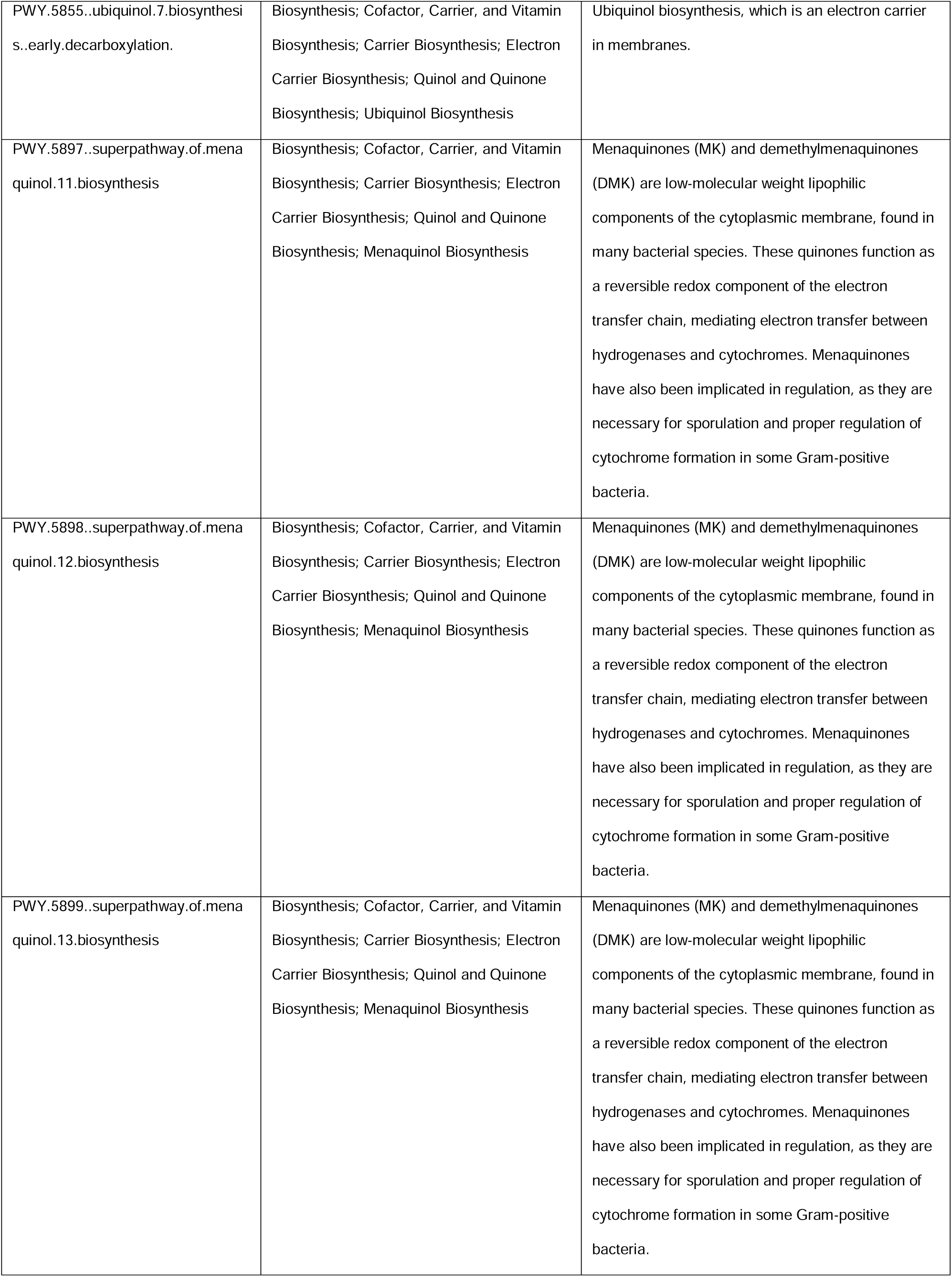
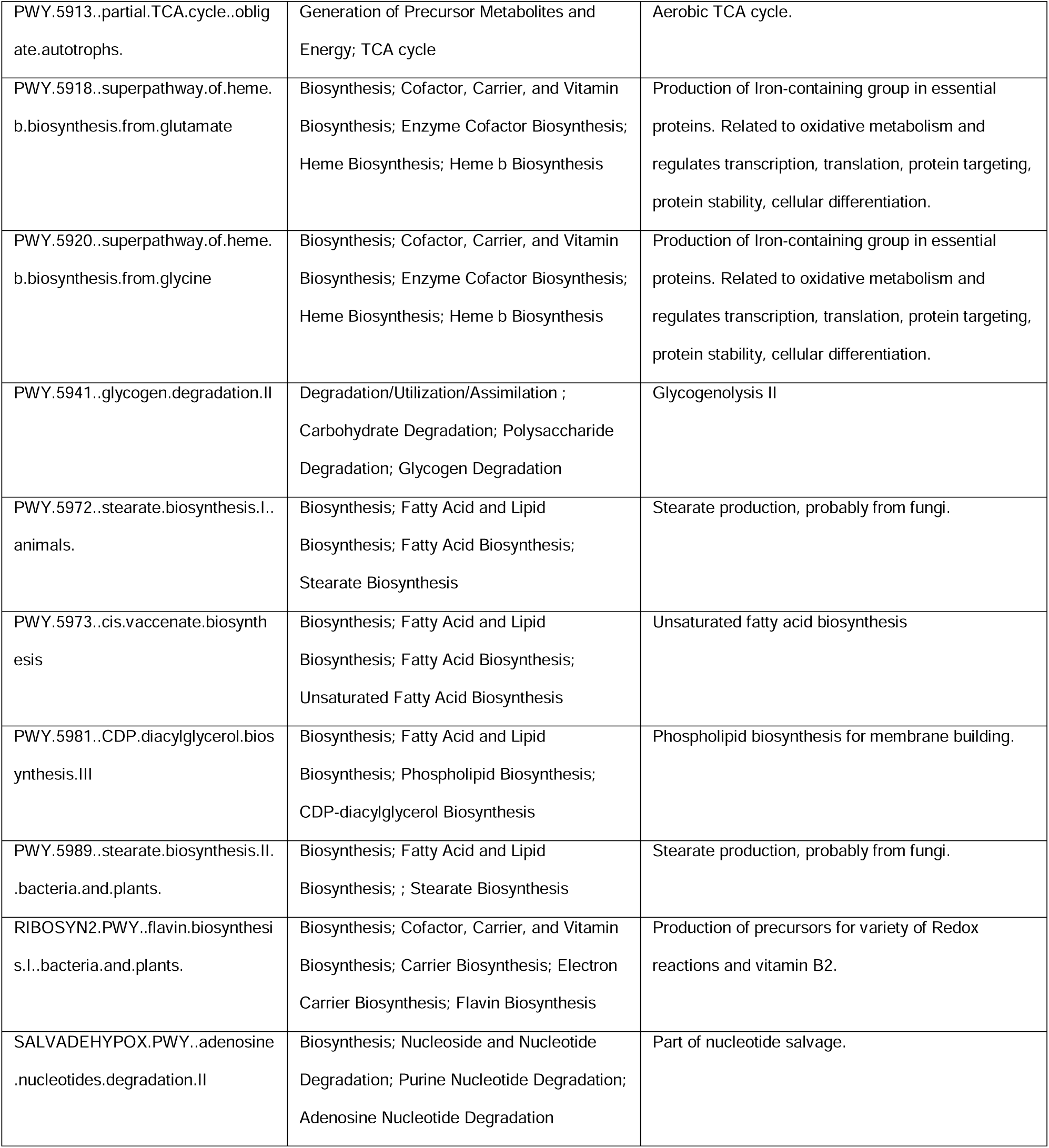
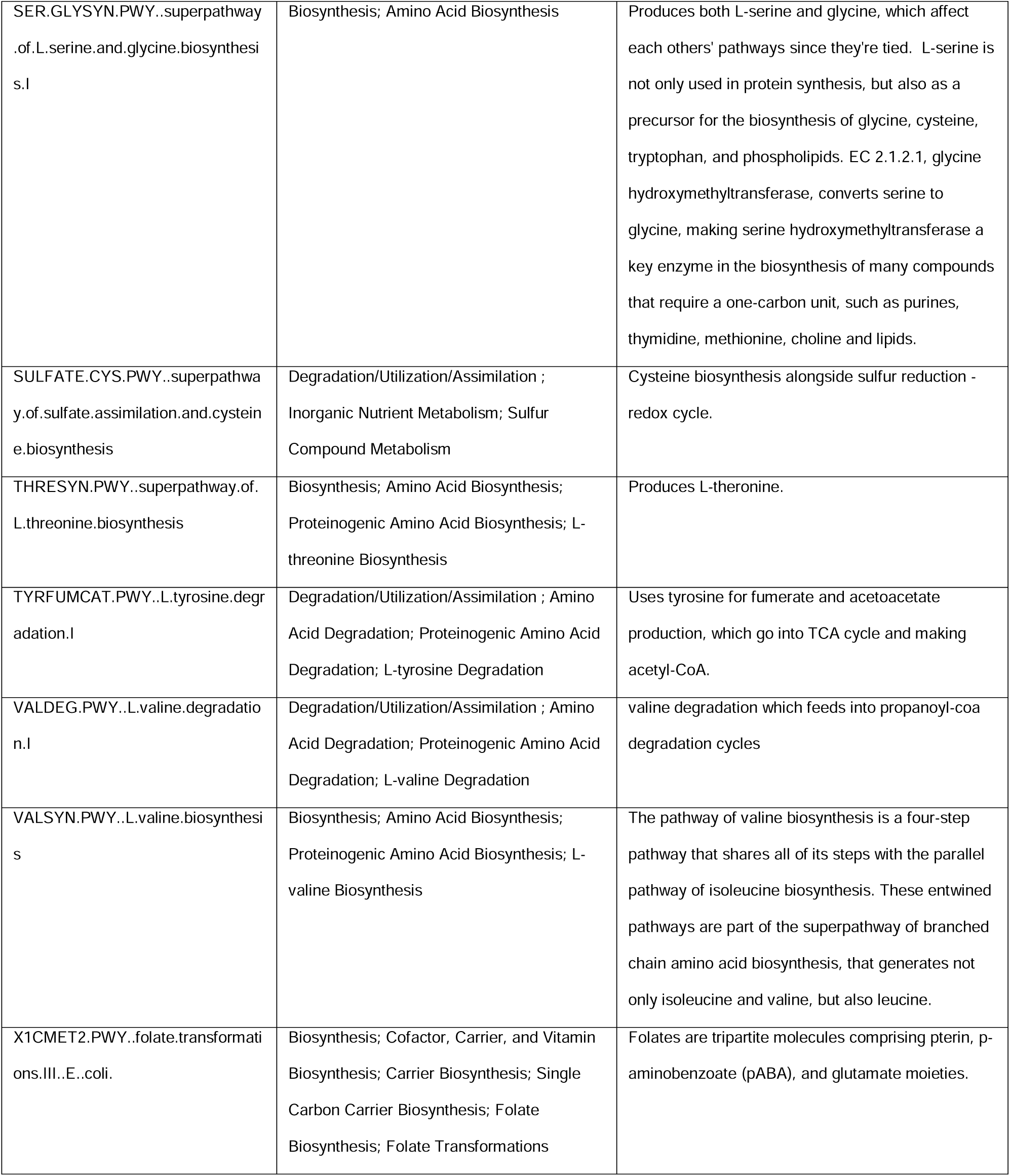
A table of the significantly associated pathways, noted using BioCyc ID, from Figure 5B, their superclasses from BioCyc, and the information we were able to find about them through BioCyc and a literature search.

